# Abnormal white matter development during early childhood in autism and developmental disability

**DOI:** 10.1101/2023.11.15.567183

**Authors:** Yuqi Liu, Jiaying Zhang, Edmund T. Rolls, Yuan Dai, Shujie Geng, Lin Deng, Zilin Chen, Yue Zhang, Minyi Tao, Lingli Zhang, Tai Ren, Jianfeng Feng, Miao Cao, Fei Li

## Abstract

The abnormal maturation of white matter microstructure in autism spectrum disorder (ASD) compared with typical developing (TD) and developmental delay/intellectual disorder (DD/ID) during early childhood and their associations with clinical implications remain unclear. Here, we characterized spatial and temporal patterns of white matter development in 364 children (including ASD, DD/ID and TD) under 8 years old using neurite orientation dispersion and density imaging. Using neurite density index (NDI), we consistently detected three white matter clusters across all the groups through clustering analysis, indicating the conserved spatial layout in atypical development. Further, three developmental stages with successively lower growth rates were identified in ASD, similar to those in TD but delayed overall. Meanwhile, DD/ID children showed the slowest growth rates and lacked staging features. Finally, for ASD children, the NDI was correlated with cognitive impairments at the fast developmental stage but associated with social deficits during the moderate-to-slow stage.

## Introduction

Autism spectrum disorder (ASD) is a typical early-childhood-onset neurodevelopmental disorder (NDD)^1^, and the presentation of its symptoms starts in early childhood. Early signs of ASD include poor skill acquisition for language and communicative gestures. Typically, characteristic ASD symptoms, including behavioural or cognitive rigidity, a lack of interest in socializing, and restricted interests, become increasingly apparent as a child develops^2^. Notably, ASD often co-occurs with developmental delay/intellectual disability (DD/ID), another typical early-childhood-onset NDD, at a high prevalence, causing additional cognitive impairments^2^. Nearly two thirds of ASD individuals younger than 4 years old exhibit comorbid DD^3^, and approximately one third of ASD individuals display comorbid ID at later ages^4^. Therefore, it is critical to identify abnormal brain developmental patterns of ASD children during the early period of life compared with typically developing (TD) and non-autism DD/ID children, to reveal the neurophysiology of the development of symptoms and cognitive impairment. Accumulating neuroimaging findings in young children indicate that the early development of white matter is highly dynamic and elaborate^5–7^. As the key feature of white matter, axons connect neurons, and their morphological development provides crucial support for the maturation of brain functions. Naturally, a series of critical questions arise: What are the specifically altered maturation patterns of axon morphology during early childhood in ASD compared to TD and DD/ID children? What are the associations between these alterations and clinical implications?

Atypical features of axon morphology in the white matter of individuals with ASD were identified by postmortem studies, exemplified by lower axon numbers and smaller axon sizes in the corpus callosum (CC)^8^. Despite providing invaluable information on brain pathology, these studies had inherent limitations, such as the scarcity of brain samples from young children, the limited sample size of postmortem brain tissue, and the absence of information on donors’ behavioural profiles. In contrast, diffusion MRI techniques provide non-invasive windows for quantifying *in vivo* white matter microstructure. In particular, using diffusion tensor imaging (DTI), previous studies found that compared to TD children, ASD and DD/ID children exhibited reduced fractional anisotropy (FA) with slower increases and flatter developmental trajectories in white matter tracts^9,10^. These findings suggest the presence of atypical white matter growth in individuals with ASD and DD/ID. However, DTI metrics lack specificity for tissue properties^11^. For example, FA can be affected by factors such as the axon packing density and the spatial coherence of axon alignments. Therefore, changes in FA may not necessarily be linked to alterations in specific axon morphological features^12^.

Neurite orientation dispersion and density imaging (NODDI)^13^ provides tissue-specific quantifications of white matter microstructure by exploiting biophysical models to disentangle the water diffusion of a voxel into each compartment. NODDI metrics, including the neurite density index (NDI) and orientation dispersion index (ODI), represent axon density and the spatial coherence of axon alignments in white matter, respectively, which were histologically validated using postmortem human neural samples^14^. Previous neurodevelopmental studies using NODDI revealed that the NDI was sensitive to developmental processes^15^. For instance, the NDI of white matter across the entire brain increased with age in TD, characterized by aninitial rapid growth phase followed by a more gradual increase^15^. Furthermore, the maturation curves of the NDI in TD exhibited spatial heterogeneity across the cerebral white matter, where callosal fibres developed earlier than in the association and projection tracts^15^. In addition, previous studies reported associations between abnormal NDI values and clinical implications in ASD adults without cognitive impairment (i.e., ASD without DD/ID)^16–19^. However, the temporal and spatial developmental patterns of axon morphological features in ASD children with different cognitive levels are still yet to be explored.

To answer these questions, we employed a large multi-shell diffusion MRI dataset comprising 364 children aged under 8 years old, including 156 children diagnosed with ASD (105 with DD/ID and 51 without DD/ID), 160 TD and 48 children diagnosed with DD/ID. We first assessed the sensitivity of NODDI in early childhood. Subsequently, we conducted hybrid hierarchical clustering to elucidate the spatial patterns of white matter development during early childhood in both typical and atypical development using NODDI. We further used the generalized additive model (GAM), a nonlinear curve fitting method, to model the developmental curves of all identified white matter clusters. We then delineated the growth rates of NODDI metrics using the first derivatives of the estimated developmental curves and subsequently defined growth stages. Finally, in these white matter clusters, we explored the group differences in NODDI metrics as well as their relationships with clinical measures in children with atypical development (Fig. 1).

**Fig. 1.**
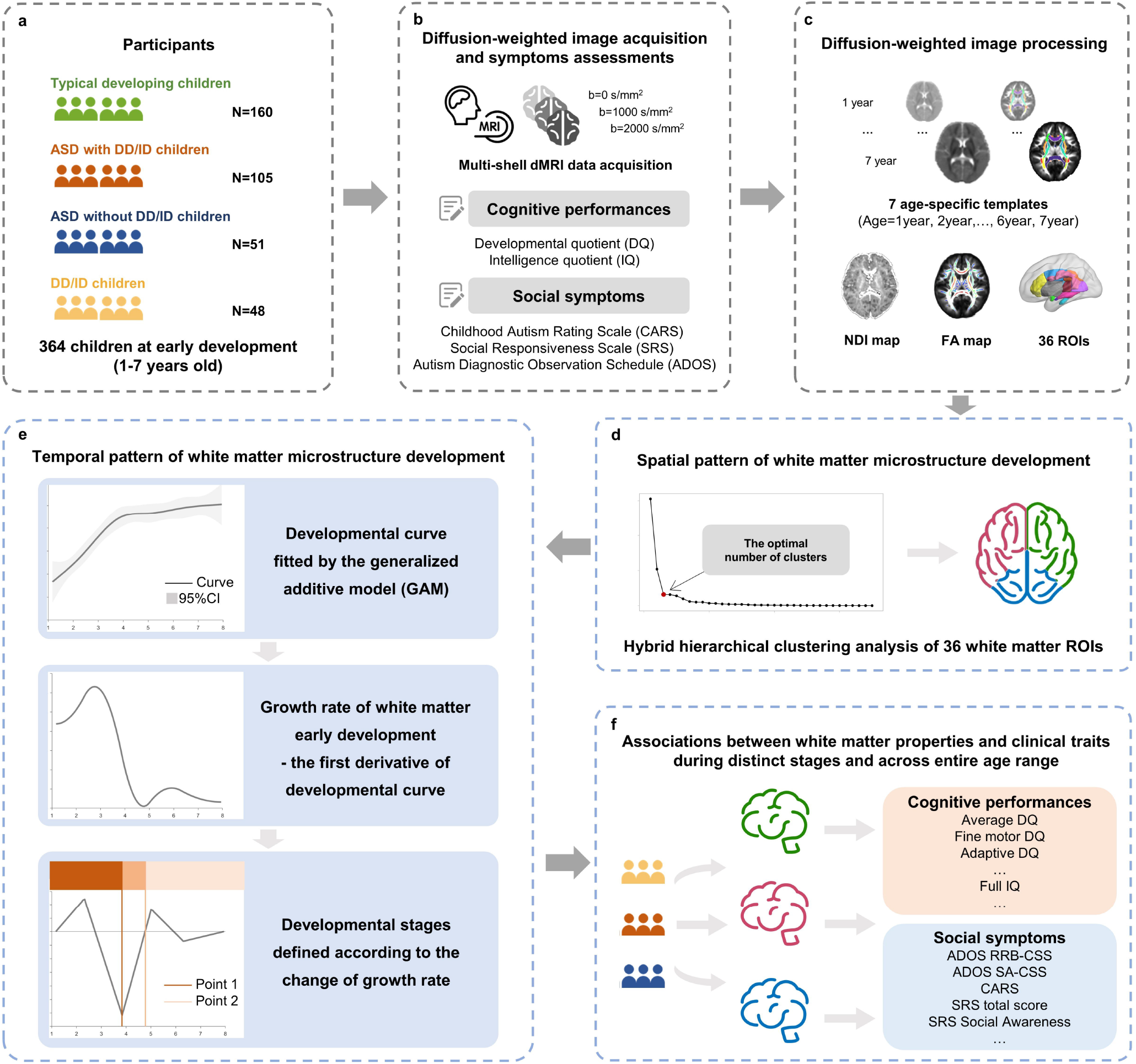
Flow diagram for the study of spatial and temporal patterns of white matter microstructure development in the ASD subgroups and DD/ID and TD groups. **a-b**, diffusion-weighted MR images and behaviour measurements were acquired from 364 children aged under 8 years old, including 156 children diagnosed with ASD (105 with DD/ID and 51 without DD/ID), 48 children diagnosed with only DD/ID, and 160 TD children. **c**, one group-specific tensor template and seven age-specific tensor templates were created for registration. In addition, DTI and NODDI were applied to compute FA and NDI maps, respectively. **d**, **a** hybrid hierarchical clustering algorithm was employed to delineate the spatial developmental patterns. **e**, temporal patterns of white matter microstructure development were characterized by growth curves, rates and developmental stages. Developmental curves were fitted for each cluster and each group. **f**, distinct associations between white matter microstructure and clinical manifestations (cognitive performance and social symptoms) of each abnormal-development group during the distinct stages and across the entire age range.

## Results

### Sample characteristics

After MR image quality control, we included a large sample of 364 participants in early childhood, comprising 160 TD children (age range: 1.17-7.94 years, 49.4% males), 51 children diagnosed as ASD without DD/ID (age range: 1.73-7.95 years, 82.4% males), 105 children diagnosed as ASD with DD/ID (age range: 1.62-7.89 years, 87.6% males), and 48 children with DD/ID (age range: 2.32-7.77 years, 35.2% males). A total of 40 participants were excluded after MR image quality control (see the Supplementary material for details about the exclusion criteria and Supplementary Fig. 1). For clinical measurements and cognitive performance, we found the following: 1) as expected, TD children had a higher average developmental quotient (DQ) and full intelligence quotient (IQ) than all three groups of children with atypical development (*P*s<1×10^-8^); 2) there was no significant difference in either the DQ or IQ between the DD/ID and ASD with DD/ID groups (*P*s>0.7); and 3) both of the ASD subgroups had more severe social deficits than the TD and DD/ID groups (*P*s<1×10^-8^). More details on the characteristics of the cohort in this study can be found in the Methods section and Supplementary Tables 1 and 2.

### Consistent clustering outcomes of white matter across early childhood in typical and atypical development

We first examined the sensitivity of NODDI metrics to white matter development across early childhood using partial Pearson correlation analyses, including sex and head motion during the MRI scan as covariates. Our results revealed that the NDI increased rapidly with age (TD: r=0.765, ASD with DD/ID: r=0.767, ASD without DD/ID: r=0.601, DD: r=0.500, all groups’ *P*s<0.001). Moreover, the ODI exhibited a plateau pattern with no age effects (TD: r=-0.030, ASD with DD/ID: r=0.111, ASD without DD/ID: r= 0.069; DD: r= –0.024, all groups’ *P*s>0.4) throughout early childhood (Supplementary Fig. 2). This observation suggested that the NDI, a marker of axon density in white matter derived from NODDI, is sensitive to the white matter developmental process in early childhood.

We next used the NDI to reveal the distinct spatial layout of white matter development during early childhood. We grouped core white matter regions into clusters using a hybrid hierarchical clustering algorithm^20^ based on the similarity of the mean NDI. A total of 36 core white matter regions were carefully defined using the ICBM-DTI-81 white matter atlas^21^ and the white matter skeleton of the study-specific group template (see Fig. 1c, Methods and Supplementary Table 3). The clustering outcomes were consistent in all four groups in early childhood: the optimal number of clusters according to the knee plots of inter-cluster Euclidean distances was three (Fig. 2a and Supplementary Fig. 3e-g for the hierarchical dendrograms for the TD, ASD without DD/ID, ASD with DD/ID and DD/ID groups, respectively; Supplementary Fig. 3a-d for plots of the searches for the optimal cluster numbers).

**Fig. 2.**
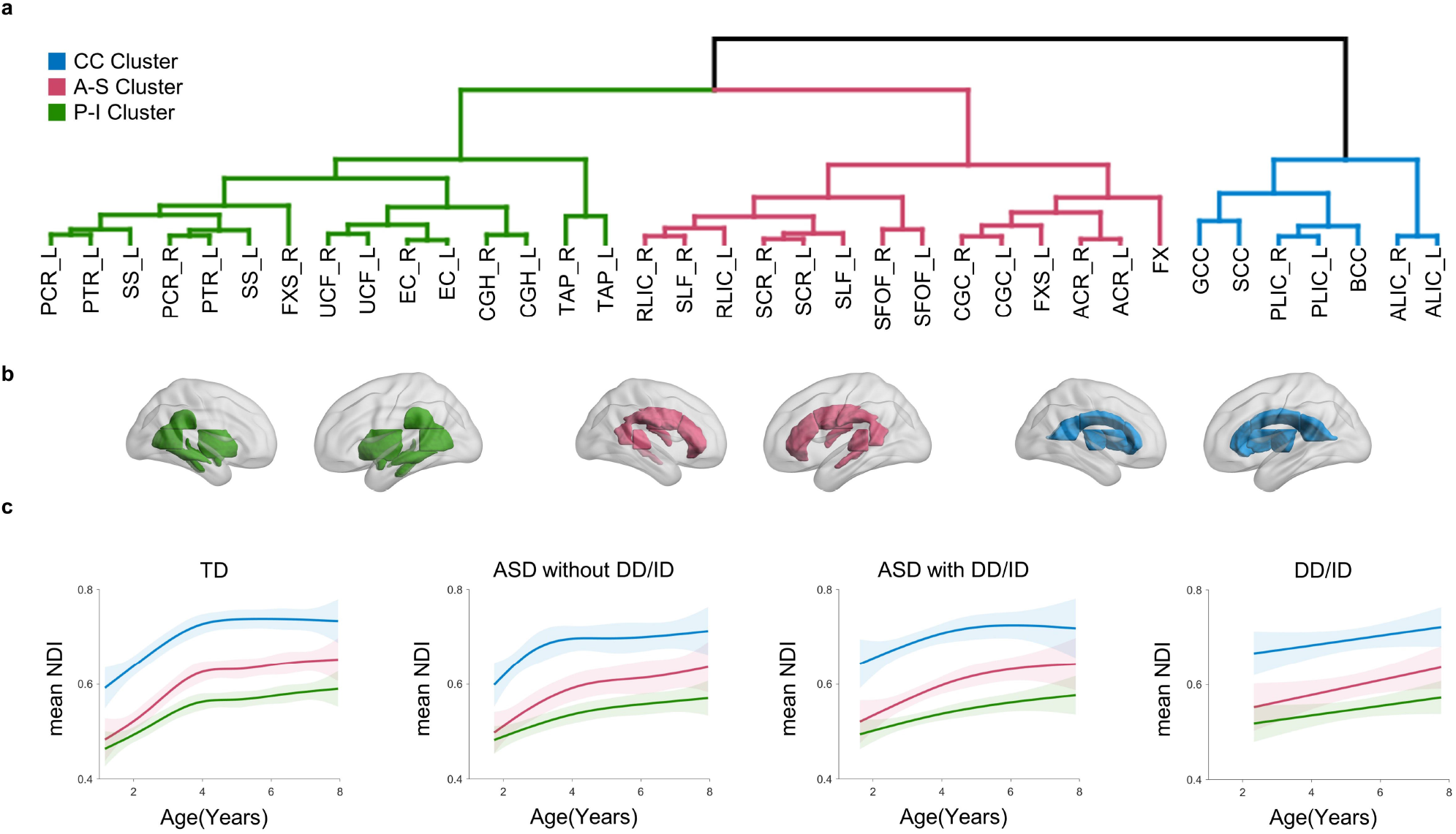
Consistent clustering outcomes of white matter regions across early childhood in children with typical and atypical development. **a**, hybrid hierarchical dendrogram in the TD group. A total of 36 cerebral white matter regions were grouped into three clusters based on the similarity of the NDI. The CC Cluster was composed of the corpus callosum (CC), anterior limb of internal capsule (ALIC) and posterior limb of internal capsule (PLIC). The anterior-superior Cluster (A-S Cluster) was composed of the anterior corona radiata (ACR), superior corona radiata (SCR), retrolenticular internal capsule (RLIC), superior longitudinal fasciculus (SLF), superior fronto-occipital fasciculus (SFOF), left fornix (cres)/stria terminalis (FX/ST-L), fornix (FX), and cingulum bundle (CGC). The P-I Cluster was composed of the posterior corona radiata (PCR), posterior thalamic radiation (PTR), sagittal stratum (SS), external capsule (EC), right fornix (cres)/stria terminalis (FX/ST-R), uncinate fasciculus (UNC), cingulum bundle (CGH), and tapetum of commissural tracts. **b**, the spatial distribution of the white matter regions of the three clusters. **c**, developmental curves of the NDI were fitted for each cluster and each group.

These three clusters of core white matter regions were as follows: 1) Cluster 1, mainly comprising the CC (blue in Fig. 2b, referred to as the CC Cluster); 2) Cluster 2, mainly composed of anterior and superior white matter (pink in Fig. 2b, referred to as the A-S Cluster); and 3) Cluster 3, mainly composed of posterior and inferior white matter (green in Fig. 2b, referred to as the P-I Cluster) (see Supplementary Table 3 for the list of white matter regions of interest (ROIs) in each cluster).

### Developmental curves of white matter clusters in early childhood

We next delineated the developmental curves of white matter clusters using the GAM^22^. For all three white matter clusters in the typical and atypical development groups, the NDI increased with age throughout early childhood (TD: *P*s<1×10^-15^; ASD without DD/ID: *P*s<1×10^-15^; ASD with DD/ID: *P*s<1×10^-5^; DD/ID: *P*s*<*0.016) (Fig. 2c). Notably, the CC Cluster always had the highest mean NDI. The P-I Cluster was characterized by the lowest mean NDI and the flattest developmental curve. Furthermore, the A-S Cluster showed the most rapid increase in the NDI in the initial period of early childhood development.

Additionally, each group exhibited unique developmental curves of white matter clusters (Fig. 2c). Specifically, the TD group exhibited the longest period of rapid increase in early childhood. The ASD without DD/ID group had a similar rapid increase but only in the initial phase of early childhood. In contrast, the ASD with DD/ID group showed a flatter increase pattern but lacked obvious differences in velocity. Moreover, the DD/ID group had the flattest developmental curves with the least increase throughout early childhood.

### Growth rates and developmental stages of early white matter maturation

For the white matter clusters, we next defined the growth rates of the NDI using the first derivatives of the developmental curves. We observed that all the groups except the DD/ID group exhibited a similar pattern of white matter growth, with a rapid increase at first and a gradual transition into a platform (solid line in Fig. 3). Moreover, the groups with DD/ID (ASD with DD/ID and DD/ID groups) always had lower growth rates across all white matter clusters than the groups without DD/ID (ASD without DD/ID and TD groups).

**Fig. 3.**
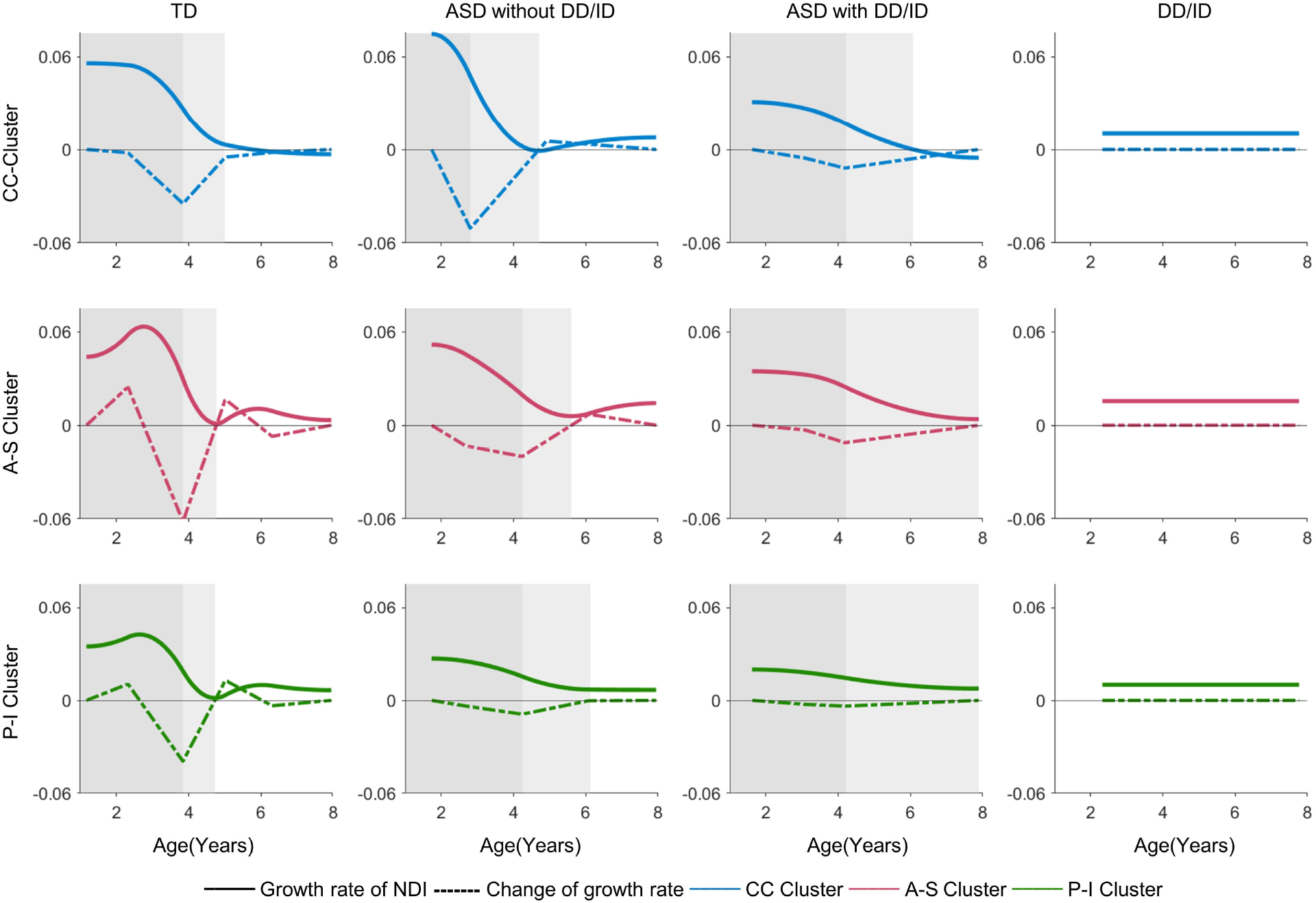
Developmental curves of white matter clusters in early childhood. The growth rates of the NDI were calculated using the first derivatives of developmental curves (solid lines) and changes in rates were represented by the derivatives of the growth rates (dotted lines). Based on the changes in rates for all clusters, three developmental stages were defined using two special timepoints: the fast, moderate and slow stages (represented by dark, grey and white colours), except for DD/ID group. Growth rates and developmental stages of the TD (column 1), ASD without DD/ID (column 2), ASD with DD/ID (column 3), and DD/ID (column 4) groups.

We then defined white matter developmental stages during early childhood according to changes of rate. Accordingly, early development was divided into three stages: a fast stage with a high rate, a moderate stage with a relatively low rate, and a slow stage with a low rate (shadowed areas in Fig. 3). Specifically, the TD group remained in the fast developmental stage until 3.84 years of age and entered the slow stage at about 5 years of age across all white matter clusters. The two ASD subgroups had different durations of the fast stages across white matter clusters. The fast stage of the A-S Cluster and P-I Cluster stopped after 4 years of age, which was slightly longer than the fast stage of the TD group. In contrast, the fast stage of the CC cluster ended before 3 years of age in the ASD without DD/ID group. Additionally, the ASD with DD/ID group entered the slow stage after 6 years of age in the CC Cluster but had almost no slow stage in the other clusters. In contrast, the DD/ID group had no clearly defined developmental stages, as it lacked variability in the growth rate of the NDI throughout early development (see Supplementary Table 4 for the time points of the developmental stages).

We next performed group comparisons of the growth rates across all four groups with normal and abnormal development (Fig. 4 middle column). During the fast stage, the growth rates of the ASD with DD/ID group were the lowest across all clusters (*P*s<1×10^-9^). In addition, the ASD without DD/ID group had significantly lower rates in the A-S Cluster and P-I Cluster but a higher rate in the CC Cluster than the TD group (*P*s<1×10^-6^). Notably, as the slow stage did not emerge in some scenarios, we combined the two non-fast stages—the moderate and slow stages—into one, a moderate-slow stage, for the following analysis. In the moderate-slow stage, the growth rates of the A-S Cluster and P-I Cluster in both ASD subgroups were significantly higher than those in the TD group (*P*s<0.004, false discovery rate (FDR)-corrected *P*s<0.01). When we examined the entire age range, we found that the ASD with DD/ID group had significantly lower growth rates in all white matter clusters than the TD and ASD without DD/ID groups (*P*s<0.02, FDR-corrected *P*s<0.05). Moreover, compared with the TD group, the ASD without DD/ID group showed a higher growth rate in the CC Cluster (*P=*0.019, FDR-corrected *P*<0.05) but a lower growth rate in the P-I Cluster (*P=*0.020, FDR-corrected *P*<0.05). In addition, the growth rates of the DD/ID group were significantly lower than those of the TD group in the P-I Cluster (*P*<1×10^-4^, FDR-corrected *P*<0.05) and lower than those of the ASD without DD/ID group in the CC Cluster and A-S Cluster (*P*s<0.03, FDR-corrected *P*s<0.05) (Supplementary Table 5).

**Fig. 4.**
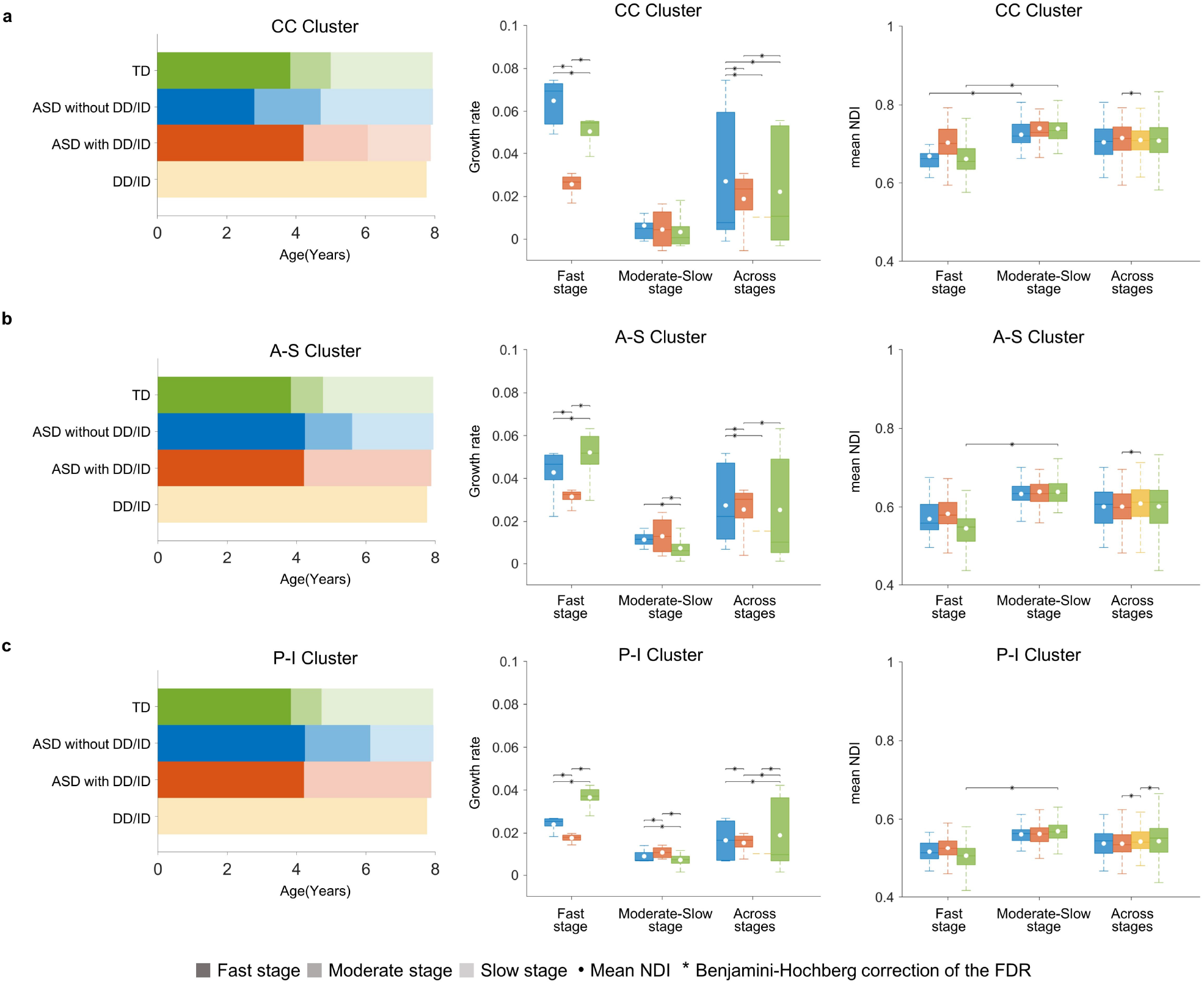
Growth rates and developmental stages of early white matter development. **a-c**, developmental stages and box plots of the group-comparison within the CC Cluster, A-S Cluster and P-I Cluster, respectively. **Left column,** developmental stages and timepoints of each stage across all groups and all clusters. **Middle and right columns**, box plots of group comparisons of the growth rates and NDI values in each stage and across the entire age range in the CC Cluster (row 1), A-S Cluster (row 2) and P-I Cluster (row 3). The moderate and slow stages were combined into the moderate-slow stage for statistical analyses.

We also performed group comparisons of the mean NDI of the white matter clusters according to the developmental stages (Fig. 4, right column). The TD group showed a significantly higher NDI in the moderate-slow stage than in the fast stage in all white matter clusters (*P*s<0.007). Meanwhile, for the ASD without DD/ID group, the NDI in the moderate-slow stage was higher than that in the fast stage in the CC Cluster (*P=*0.033). At each developmental stage, there were no significant group differences in the NDI for any of the white matter clusters. Across the entire age range, the DD/ID group had a lower NDI than the TD group in the A-S Cluster (*P=*0.014, FDR-corrected *P*<0.05) and a lower NDI than the ASD with DD/ID group across all clusters (*P*s<0.003, FDR-corrected *P*s<0.05), while there was no significant difference in the NDI among the two ASD subgroups and TD group (Supplementary Tables 6 and 7).

### Associations between the NDI of white matter clusters and social symptoms and cognitive performance in ASD and DD/ID children

To evaluate the clinical relevance of the neuroimaging findings, we next performed partial correlations between the mean NDI of white matter clusters and the social symptoms and cognitive performance in young children with abnormal development (including the ASD without DD/ID, ASD with DD/ID and DD/ID groups). The relationships between the NDI of white matter clusters and clinical measurements of the ASD subgroups varied across developmental stages. In the fast stage, for the ASD with DD/ID group, we observed the following: 1) the NDI in the P-I Cluster was negatively correlated with the adaptive DQ (r=-0.343, *P*=0.006, FDR-corrected *P*<0.05), average DQ (r=-0.313, *P*=0.012, FDR-corrected *P*<0.05) and fine motor DQ (r=-0.365, *P*=0.031); 2) the NDI in the A-S Cluster was negatively correlated with the fine motor DQ (r=-0.443, *P*=0.008, FDR-corrected *P*<0.05), adaptive DQ (r=-0.252, *P*=0.045), and average DQ (r=-0.298, *P*=0.017) (Fig. 5a; Supplementary Table 8a and b). Notably, the moderate and slow developmental stages were combined into the moderate-slow stage due to their short durations (the same reason mentioned in the previous subsection). At the moderate-slow stage, for the ASD without DD/ID group, the NDI of the CC Cluster was negatively correlated with the scores of social deficits measured by the Social Awareness subscale of the Social Responsiveness Scale (SRS)^23^ (r=-0.565, *P*=0.006, FDR-corrected *P*<0.05) (Fig. 5b; Supplementary Table 8c and d). Across the entire age range, we identified that for the ASD without DD/ID group, the NDI in the CC Cluster was negatively correlated with Autism Diagnostic Observation Schedule (ADOS)^24^ restricted, repetitive behaviour calibrated severity score (RRB_CSS) (r=-0.370, *P*=0.011, FDR-corrected *P*<0.05). In the DD/ID group, there was no significant correlation between the NDI and clinical measurements (Fig. 5b; Supplementary Table 9a-c).

**Fig. 5.**
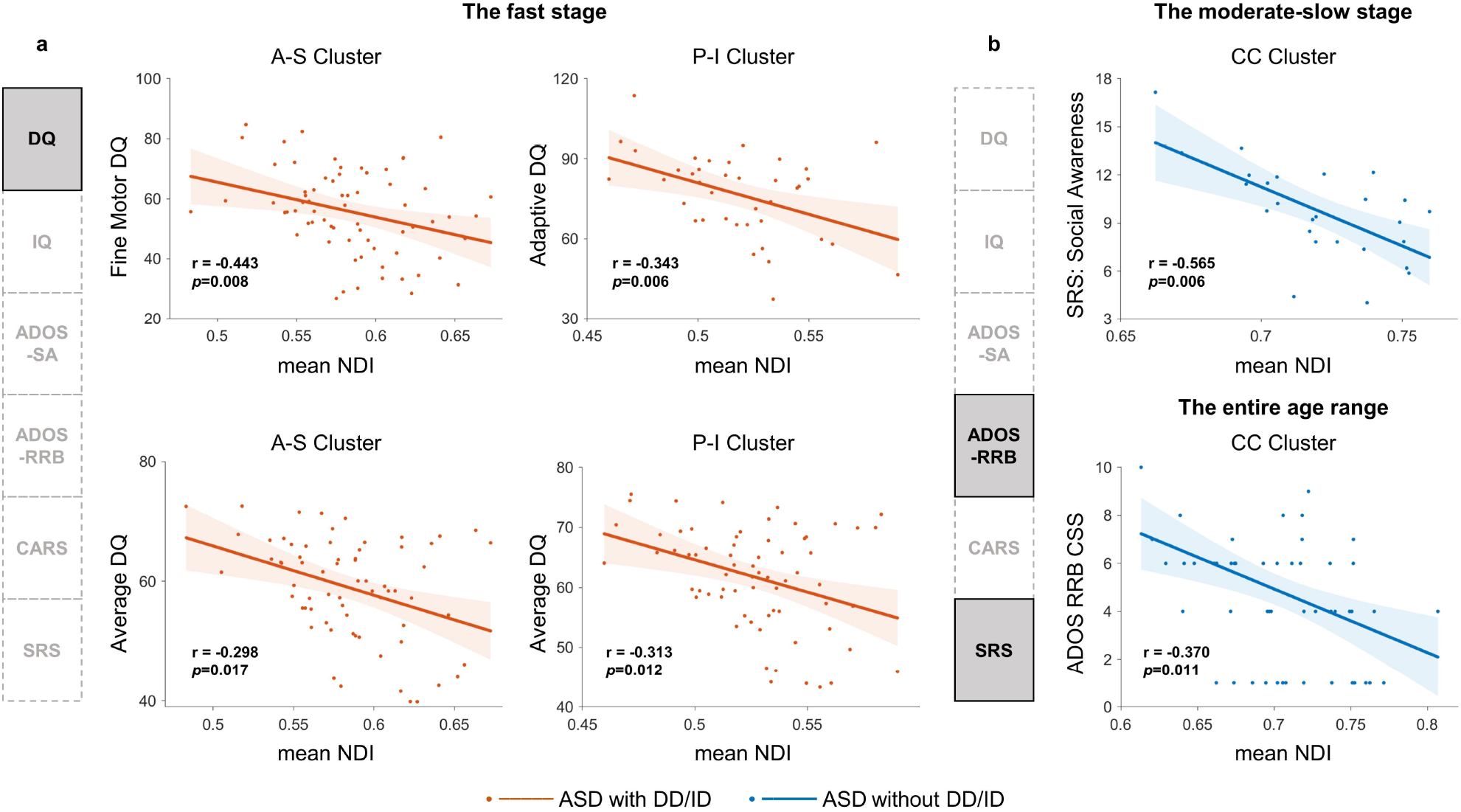
Associations between the NDI of white matter clusters and social symptoms and cognitive performance in children with NDDs. **a**, during the fast stage, the ASD with DD/ID group showed negative correlations between the NDI and DQ (measuring cognitive capacity and assessed by the Gesell Developmental Schedules (GDS)). The NDI in the A-S cluster was negatively correlated with the fine motor DQ (r=-0.443, *p*=0.008) and average DQ (r=-0.298, *p*=0.017) (left column 1). The NDI in the P-I cluster was negatively correlated with the adaptive DQ (r=-0.343, *p*=0.006) and average DQ (r=-0.313, *p*=0.012) (left column 2). **b**, During the moderate-slow stage, the NDI of the CC cluster in the ASD without DD/ID group was negatively correlated with the scores of social deficits measured by the Social Awareness of Social Responsiveness Scale (SRS) (r=-0.565, *p*=0.006) (the upper-right box). Across the entire age range, the ASD without DD/ID group showed a negative correlation between the NDI in the CC cluster and the restricted, repetitive behaviour calibrated severity score (RRB_CSS) of the Autism Diagnostic Observation Schedule (ADOS) (r=-0.370, *p*=0.011) (the lower-right panel). The intelligence quotient (IQ) was measured by the Wechsler Preschool and Primary Scale of Intelligence (WPPSI) and Wechsler Intelligence Scale for Children-Revised (WISC-R). CARS, Childhood Autism Rating Scale.

### Validation analyses

To evaluate the reliability of our findings, we performed a split-half validation analysis. Due to the limited sample sizes in the ASD without DD/ID and DD/ID groups, the validation analyses were only performed in the ASD with DD/ID and TD groups. Specifically, we divided 105 ASD with DD/ID and 160 TD children into two validation populations: 1) Validation Population 1, 53 ASD with DD/ID children (7 females) and 80 TD children (41 females); 2) Validation Population 2, 52 ASD with DD/ID children (6 females) and 80 TD children (39 females). We applied the hybrid hierarchical clustering algorithm and curve fitting to these four validation groups separately. For the clustering analyses, most results of the validation populations were consistent with the main findings except for those of 2 white matter regions (right fornix (cres) and/or stria terminalis and left tapetum). In the TD group of Validation Population 1, the right fornix (cres) and/or stria terminalis was clustered into the A-S Cluster, which belonged to the P-I Cluster in the main clustering findings. In the TD and ASD with DD/ID groups of Validation Population 2, the left tapetum was clustered into the A-S Cluster instead of the P-I Cluster in the main clustering findings. Given that our analyses in the main findings included 36 regions in total, alterations in only 2 white matter regions in the clustering outcomes were still acceptable. Moreover, the developmental curves for all validation populations were almost consistent with the main results (Supplementary Fig. 4a and b). The only exception was that the developmental curve of the P-I Cluster of ASD with DD/ID children in Validation Group 1 tended to be linear, suggesting a stronger similarity with the DD/ID population.

In addition, we conducted another validation analysis with male-only samples for all groups to explore the potential sex effects of ASD. A total of 250 male participants were included (79 TD, 37 DD/ID, 42 ASD without DD/ID, and 92 ASD with DD/ID). We applied the hybrid hierarchical clustering algorithm and curve fitting to all four male-only groups. For clustering analyses and developmental curves, the results across the four groups were also mostly consistent with the main findings. A minor change was observed in the developmental curve of the A-S Cluster for the DD/ID group, whose growth rates were still very slow but showed mild staging features (Supplementary Fig. 5a and b).

## Discussion

Using NODDI, we investigated the spatial and temporal patterns of white matter microstructure development and their relationship with clinical traits in two ASD subgroups compared with TD and DD/ID groups in early childhood. This study yielded three primary insights. First, consistent with the TD group, the same three white matter clusters were identified among the two ASD groups and the DD/ID group, indicating that the spatial layout of white matter was conserved in children with both typical and atypical early development. Second, three stages with successively lower developmental rates were identified in children with ASD, which were similar to those of TD children but delayed overall with a more severe manifestation in ASD with DD/ID children. In contrast, the DD/ID group was characterized by uniform and slow rates without defined stages. Last, the cognitive performance of the ASD with DD/ID group were correlated with the NDI in the A-S and P-I clusters in the fast developmental stage, while the social deficits of the ASD without DD/ID group were correlated with the NDI of the CC Cluster during the moderate-slow stage.

In this study, we observed the unique spatial layout of axon morphological features in white matter during early typical and atypical development. Although previous studies indicated spatial heterogeneities in white matter during development^15^, we classified the cerebral white matter regions into three clusters based on the early developmental pattern of axon density for the first time. Our findings suggested that axon density was not spatially evenly distributed, and various biological mechanisms in distinct regions may underlie the spatial developmental pattern during early childhood, including axon growth and axon alignments in white matter^25^. Specifically, the CC Cluster, mainly including the CC, was characterized by the highest NDI value among the three clusters over the whole age range studied. Previous studies revealed that the CC exhibited rapid growth in the first 2 years with the earliest maturation^15^, which might lead to the high density of axon packing during early childhood. Moreover, the CC is the major white matter tract for interhemispheric functional communications^26^. The high demand for such communications may be a key driving factor of the high axon density in this white matter fibre bundle during early childhood. In contrast, the A-S Cluster exhibited the highest growth rates among the three clusters. White matter in the A-S Cluster is mainly intra-hemispheric and projects to the frontal, parietal and temporal cortices. Their fast growth would foster efficient and coordinated transmissions of signals across the entire brain^27^ and may facilitate rapid skill acquisition, including language and social interaction skills, in the first 7 years of life^28^. In addition, we found that the A-S Cluster showed fast development until approximately 4 years of age. However, previous studies found that the most rapid growth in surface area of the lateral frontal, lateral parietal and temporal cortices takes place before 2 years old^29^. This inconsistency might be because these cortical areas may develop earlier than the core white matter. In addition, the P-I Cluster exhibited the lowest NDI value and growth rates, consistent with previous findings of the late NDI maturation until adulthood in this white matter cluster^15^. Notably, we applied GAM to fit the developmental curves of FA (a common metric derived from DTI), and found FA in white matter clusters exhibited similar patterns as previous studies^7,30^ (Supplementary Fig. 6a and b). Furthermore, the consistency of spatial developmental patterns across all four groups indicated that the spatial patterns of white matter axon density were unaffected by ASD or DD/ID during early childhood. In neurodevelopment, as the proto-map of the cytoarchitectonic areas is provided by the proliferative units in the cerebral ventricle^31^, the spatial patterns of both neurons and axons in the brain may already be finalized before the neurophysiologies of ASD emerge^32^ and remain unaffected by the expression of ASD genes afterwards. However, these questions have yet to be answered, and future studies that mimic the cellular development of the brain using organoids may be beneficial to elucidate the mechanism behind this phenomenon.

Another important finding in this study was the distinct developmental stages defined by changes in growth rates during early childhood. We detected the fast stage before 2.7 years of age with the fastest growth rates in the TD group, indicating rapid axon development before this time point. This is in line with postmortem and *in vivo* MR imaging findings that suggested a rapid increase in axon packing across the first 2-3 years of life^33,34^. The growth rates slowed during the ages of 4 to 5 years, and the axon development in the CC Cluster almost approached maturation after the age of 5 years, indicating that other factors, for example, axon diameter, might drive white matter development during late childhood. Compared to the TD group, both ASD subgroups exhibited developmental delays with reduced growth rates. Specifically, the ASD with DD/ID group showed more severe delays than the ASD without DD/ID group, possibly due to the genetic abnormalities that result in a comprehensive alteration in axonal development in individuals with DD/ID^35^. Additionally, both ASD subgroups exhibited atypical increases during the fast developmental stage, indicating overgrowth of the brain during early life in individuals with ASD^36^. Interestingly, we found that of all three clusters, the ODI, a marker of the coherence of axon alignment derived from NODDI, was within the normal range in the groups with both typical and atypical neurodevelopment and showed no significant intergroup differences or age effects during early development, consistent with previous findings that the ODI was not significantly associated with age in most white matter tracts in TD^15^. This finding suggested that in NDDs such as ASD and DD/ID, the abnormalities of core white matter during early development are associated with axon density but not the dispersion of axon alignment.

Finally, our findings unveiled the potential neurophysiological mechanisms underlying the temporal development of symptoms in children with ASD. Previous studies of ASD adults without cognitive impairment reported associations between the NDI values and clinical implications, including abnormal facial emotional recognition, visual processing and autistic symptoms^16–19^. Our findings highlighted the importance of the early childhood in ASD by revealing that symptoms of cognitive and social capabilities in children with ASD were related to axon density in white matter clusters at different developmental stages. Specifically, in the fast stage, atypical axon density growth in the A-S and P-I clusters was associated with poorer cognitive performances in the ASD with DD/ID group, especially for the fine motor DQ and adaptive DQ. This could be attributed to the associations of white matter in these two clusters with general cognitive performance, especially executive function and visual-motor performance, which would have an impact on the fine motor DQ and adaptive DQ^37,38^. In the later moderate-slow stages, axon density growth of the CC cluster was associated with milder social deficits in the ASD without DD/ID group. This further supported the importance of white matter in CC cluster in social deficits of ASD children^9^. These distinct associations between white matter clusters with behavioural domains at different developmental stages may provide evidence for behavioural evolutions in early development in ASD children, and offer invaluable insights for ASD subtyping and behavioural interventions from a developmental perspective.

Several limitations in the current study should be stated. First, though no children with ASD or DD/ID with any medical history before the MRI scan were included, some had already received behavioural interventions, especially older children, which might partly have an impact on the estimation of their representative developmental curves. Studies in neonates or even fetuses with high risk of atypical development in future could provide corroborative evidence. Currently, we are working on extending our ongoing SAED project by including participants from those populations. Second, the effect of sex is an important factor in ASD research. Although we observed consistent findings in the male-only subsample for validation, including more female participants in future studies would be helpful to reveal sex differences in the developmental patterns of children with ASD. Third, due to the challenges of collecting imaging data from children in the early childhood, our study had relatively limited sample sizes for certain groups. However, we have validated our findings within split-half validation groups, ensuring consistent results. Data of children in the early childhood with larger sample size was supposed to collect in the future and validate our findings (e.g., the ongoing data collection in the SAED cohort). Finally, we conducted this study using a cross-sectional dataset, and future studies with a longitudinal design would be beneficial to further elucidate the patterns of white matter development in this critical period of life.

In conclusion, this study revealed spatially and temporally uneven white matter microstructure development in children with ASD and DD/ID. While the ASD groups maintained the spatial configuration of axon density in core white matter, both ASD children with and without DD/ID exhibited altered temporal patterns of white matter development. In contrast, children with DD/ID were characterized by uniform and slow growth rates without staging features. Furthermore, distinct associations between white matter microstructure and clinical manifestations in the developmental stages may elucidate the targets of both key time windows and featured clinical symptoms for early behavioural interventions in children with ASD.

## Methods and Materials

### Participants

Participants were recruited in the neurodevelopmental project – Shanghai Autism Early Development Cohort (SAED)^39^ in Xinhua Hospital affiliated to Shanghai Jiao Tong University. This study included 364 children in early childhood, comprising 156 children diagnosed as ASD (105 with DD/ID and 51 without DD/ID), 48 children with DD/ID and 160 age-matched TD children. Clinical diagnoses were made by experienced clinicians based on the Diagnostic and Statistical Manual of Mental Disorders-Fifth Edition (DSM-5)^1^ and were confirmed by the ADOS^24^. The ADOS calibrated severity scores, including the social affect calibrated severity score (SA-CSS) and the RRB-CSS, were calculated to allow the comparison of autism severity across participants tested with different ADOS modules^40^. The DQ was assessed by the Gesell Developmental Schedules (GDS)^41^ for children younger than 4 years. The IQ was assessed by the Wechsler Preschool and Primary Scale of Intelligence (WPPSI)^42^ for 4-to 6-year-old children and the Wechsler Intelligence Scale for Children-Revised (WISC-R)^43^ for children older than 6 years. The Childhood Autism Rating Scale (CARS)^44^ and SRS^23^ were used to assess social-related clinical phenotypes. All participants had no neurological issues, such as suspected vision problems, hearing problems, known genetic disorders or neurological conditions (see the Supplementary methods for more details of the ascertainment and the inclusion criteria). Informed consent was obtained from the parents or the guardians of each participant. The study was conducted in accordance with the Declaration of Helsinki, with approval from the Ethical Committee of the Xinhua Hospital Affiliated to Shanghai Jiao Tong University, and in compliance with all applicable laws and regulations. All necessary biosecurity and institutional safety protocols were followed during the study.

### Multi-shell diffusion MRI acquisition

MRI scans were performed on a 3.0T Siemens Verio MRI scanner with a 32-channel head coil in the Shanghai International Medical Centre. An echo-planar imaging (EPI) sequence was employed for multi-shell diffusion-weighted MR imaging, and its parameters were as follows: 65 axial slices; echo time (TE) = 84 ms; repetition time (TR) = 14500 ms; field of view (FOV) = 192 mm × 192 mm; voxel size = 2×2×2 mm^3^; two shells with diffusion weighting of b=1000 and 2000 s/mm^2^; 30 directions for each shell; 3 additional b0 images without diffusion weighting (b = 0 s/mm^2^); a posterior-to-anterior phase-encoding direction; and a total acquisition time ∼ 15 minutes. For children with high compliance, MR scans were performed without additional preparations. Otherwise, sedation with chloral hydrate orally or by enema before the MR scan, a common preparation strategy for paediatric MR examinations, was given under the supervision and careful guidance of experienced nurses.

### Diffusion MR image quality control and processing

The image quality of diffusion MR images was visually checked by two researchers who were blinded to the diagnosis (Dr. J Zhang and M.D. Y Liu). After image quality control, diffusion-weighted MR images were first preprocessed using MRtrix3 (www.mrtrix.org) and FSL version 6.0.6 (https://fsl.fmrib.ox.ac.uk/fsl/fslwiki/FSL). The preprocessing steps were as follows: 1) denoising^45^; 2) removing the Gibbs ring^46^; 3) eddy-current distortion and motion correction using the FSL eddy^47^; and 4) B1 bias correction using the Ants algorithm^48^. After preprocessing, we computed DTI and NODDI metrics in the native space using FSL and in-house scripts with the NODDI MATLAB toolbox (https://www.nitrc.org/projects/noddi_toolbox), respectively. To minimize the potential bias that may be introduced by inaccurate registrations in early childhood, we defined the core white matter ROI masks using the following multi-level computational framework: 1) we first warped the labels of ICBM-DTI-81 white matter atlas^21^ (representing the core white matter ROIs across the brain) into the study-specific group tensor template; 2) we then multiplied the masks of the core white matter ROIs and the white matter skeleton of the whole sample; and 3) finally, we warped the labels of the skeletonized core white matter ROIs into study-specific age-specific tensor templates (one per year, seven in total). Due to incomplete coverage of the cerebellum, we chose only 36 core white matter ROIs in the cerebrum (see Supplementary Table 3). The study-specific age-specific tensor templates were created from tensor maps of a selective subset of participants within the same year (see the Supplementary Methods and Supplementary Table 10). Then, the group-specific tensor template was created using these seven age-specific tensor templates and was further used to create the white matter skeleton of the whole sample. The registrations involved were conducted using NiftiReg^49^ and DTI-TK^50^. Notably, we chose skeletonized core white matter ROIs here to 1) minimize the potential bias of inaccuracies in registration and 2) avoid the influences of subjective experience on white matter anatomy. For all the participants, their NDI maps were transformed from the native space into the space of the corresponding study-specific age-specific tensor template. The transformation matrix was estimated by registering the tensor maps from the native space into the study-specific, age-specific tensor template. Finally, for all the participants, we computed the mean NDI value of the 36 core white matter ROIs for the following statistical analyses.

## Statistical analysis

### Clustering analyses

To reveal the spatial pattern of early white matter development, we conducted hierarchical clustering analysis based on the mean NDI value of the 36 core white matter regions into clusters for all four groups. Here, the hybrid hierarchical clustering algorithm^20^ included three steps: first, bottom-up clustering was used to identify the mutual clusters, followed by the application of constrained top-down clustering in which mutual clusters could not be divided and re-application within each mutual cluster (implemented in the “hybridHclust” package in R version 3.6.0). For each group, the optimal number of clusters was chosen as the “knee point” on the inter-cluster distance vs. the number of cluster plots.

### Modelling developmental curves of white matter clusters and calculating their growth rates in the population

Inspired by the nonlinear nature of white matter development^6^, we modelled developmental curves of the NDI in each white matter cluster using a nonparametric regression approach — GAM^22^ (using the “mgcv” package in R version 4.2.1). Here, we included sex and head motion during the MR scan in the GAM. GAM can be represented by the following formula: **y_i_(age)∼**β**_0_ + s(age) +** β**_1_ *sex +** β**_2_ * AMzscore +** β**_3_ * RMzscore +r** where *yi(age)* stands for the NDI value in each white matter cluster of the i^th^ subject at the age at the time of the scan and *sex* represents the sex information of the i^th^ subject. Head motion was quantified by calculating the z score of the root-mean-square (RMS) displacement of both the mean absolute (*AMzscore*) and mean relative (*RMzscore*) intervolume displacement. *r* represents the residuals, and the nonparametric term s(*age)* represents the age of the i^th^ subject. The estimated nonparametric functions *s(age)* were obtained using cubic splines; – formula of *s(age)* was described in the Supplemental methods, and the smoothing parameter, effective degrees of freedom (EDFs), R-squared and Akaike information criterion (AIC) were shown in Supplementary Table 11. The EDF values formed by GAM showed the degree of the curvature of the relationship. Next, we employed the first derivatives of the developmental curves to represent the growth rate of the population during early childhood^6^.

### Defining stages of early white matter development

The stages of early white matter development were defined using special time points of the derivatives of the growth rates. In our scenario, two time points were identified: 1) time point 1, at which the derivative of the growth rate reached the minimum value, representing the maximal decrease in the rate of growth; and 2) time point 2, at which the derivative of the growth rate reached zero, representing the beginning of a platform period in early childhood. Notably, for the CC Cluster, there were two special cases of defining the second special point: The first was that in the TD group, the second special time point was defined at the changing point of the derivative of the growth rate at 5 years of age, which was close to zero. The second was that in the ASD with DD/ID group, the second special time point was defined as approximately 6.07 years of age when the growth rate reached 0. Therefore, for early white matter development, we defined the fast stage as the period before time point 1, the slow stage as the period after time point 2, and the moderate stage as the period between these two special time points.

### Group comparisons at various stages of early white matter development

For each white matter cluster, we performed a general linear model (GLM) in Matlab 2020a to evaluate group differences in both the NDI and its rate of growth at various stages of early development. Notably, the moderate and slow developmental stages were combined as a moderate-slow stage due to their short durations. Age, sex, and head motion during the MRI scan were included as covariates to account for potential confounding effects.

### Correlation analyses of abnormal development between the NDI of white matter clusters and cognitive performance and clinical scores

For the three groups with abnormal development, we performed correlation analyses between the NDI of white matter clusters and cognitive performance as well as clinical scores in Matlab 2020a: partial Pearson correlation for continuous variables (CARS total score; SRS, DQ and IQ scores) and partial Spearman correlation for categorical variables (CARS subscale scores; ADOS SA-CSS and RRB-CSS). For all the correlation analyses, we included age, sex, and head motion during the MRI scan as covariates. For both ASD subgroups, we conducted correlation analyses not only for the fast and moderate-slow stages but also across the entire age range. For the DD/ID group, correlation analysis was only conducted across the entire age range due to its lack of change in the growth rate of the NDI. In each group, FDR correction using Benjamini_Hochberg correction was conducted for each scale within each cluster and each stage.

## Acknowledgements

We thanked Dr. Yong He from Beijing Normal University and Dr. Daniel Alexander from University College London for their helpful comments and discussions. This study was supported by grants from the National Natural Science Foundation of China (82125032, 81930095, 81901826, 81761128035, 82202243, 82204048 and 82001771), the Science and Technology Commission of Shanghai Municipality (19410713500 and 2018SHZDZX01), the Shanghai Municipal Commission of Health and Family Planning (GWV-10.1-XK07, 2020CXJQ01, 2018YJRC03, 20214Y0125), the Shanghai Clinical Key Subject Construction Project (shslczdzk02902), the Shanghai Municipal Science and Technology Major Project [No.2018SHZDZX01], Innovative research team of high-level local universities in Shanghai (SHSMU-ZDCX20211100), the Guangdong Key Project (2018B030335001).

## Data availability

Access to the deidentified participant research data must be approved by the research ethics board on a case-by-case basis, please contact the corresponding authors for assistance in data access request.

## Author contributions

F.L., M.C. and J.Z. designed the study. Y.L., Y.D., S.G., L.D., Z.C., Y.Z., M.T., L.Z., T.R. contributed to the acquisition of research data. Y.L., J.Z. and M.C. conducted the data analysis. F.L., M.C., J.Z. and Y.L. provided the interpretation of results. M.C., J.Z. and Y.L. wrote the first draft of the manuscript. E.T.R. and J.F. revised the manuscript.

