## Supplementary Materials for "Abnormal white matter development during early childhood in autism and developmental disability"

**Supplementary Fig. 1. Diffusion-weighted MR images with the incomplete coverage of cerebral regions (occipital/parietal/temporal/frontal lobes), signal dropout, zig-zag pattern and zipper artefact.**

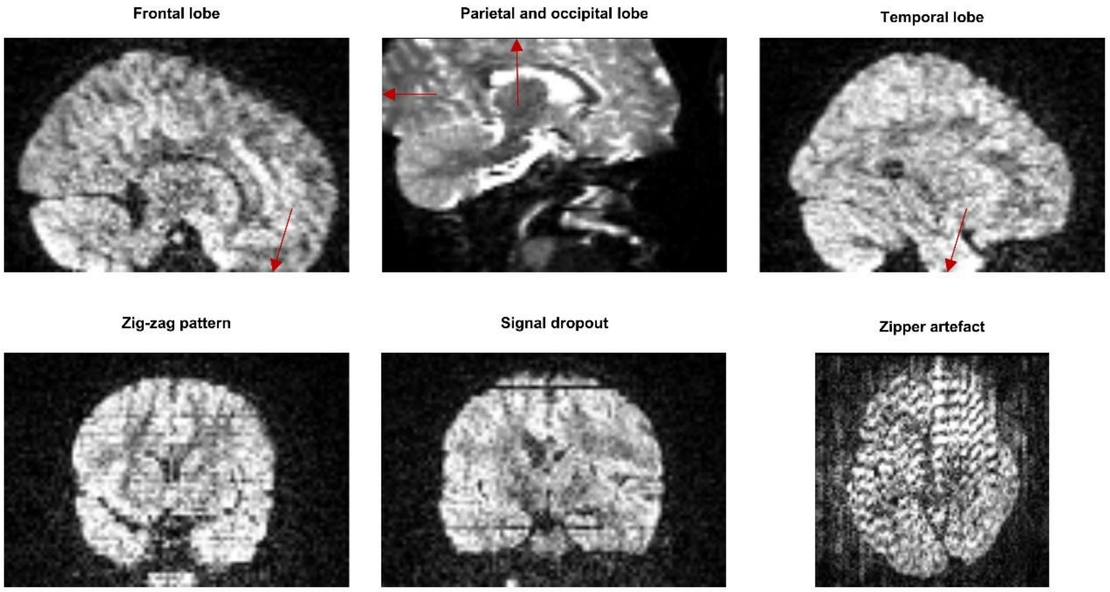

**Supplementary Fig. 2. The evolution of mean NODDI metrics across 36 white matter regions across the early childhood.** ASD, autism spectrum disorder. DD/ID, developmental delay/intellectual disability. TD, typical developing. ODI, orientation dispersion index. NDI, neurite density index.

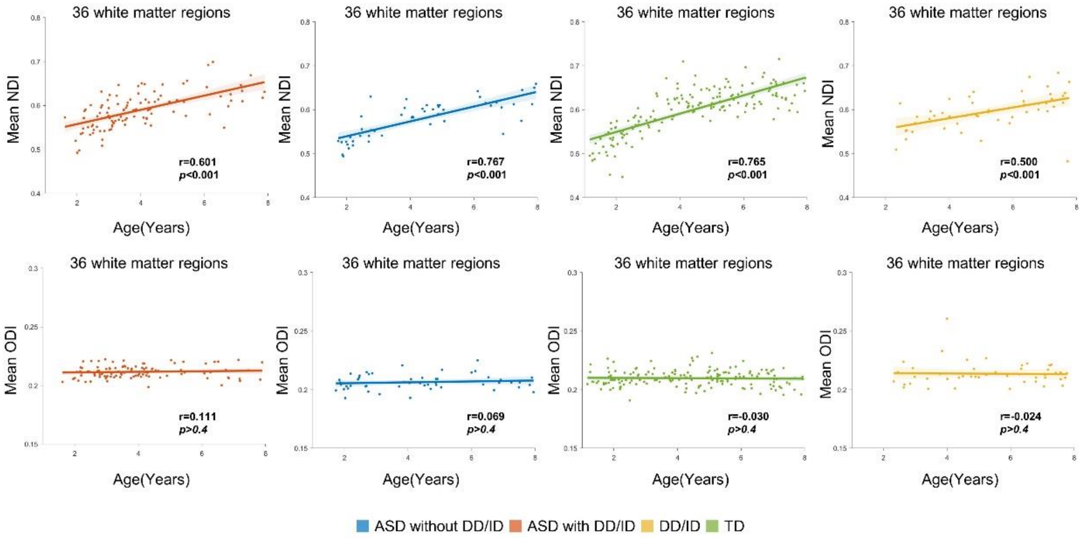

**Supplementary Fig. 3. Knee plots of inter-cluster Euclidean distances of four groups and hierarchical dendrograms of two ASD subgroups and DD/ID.** ASD, autism spectrum disorder. DD/ID, developmental delay/intellectual disability. TD, typical developing.

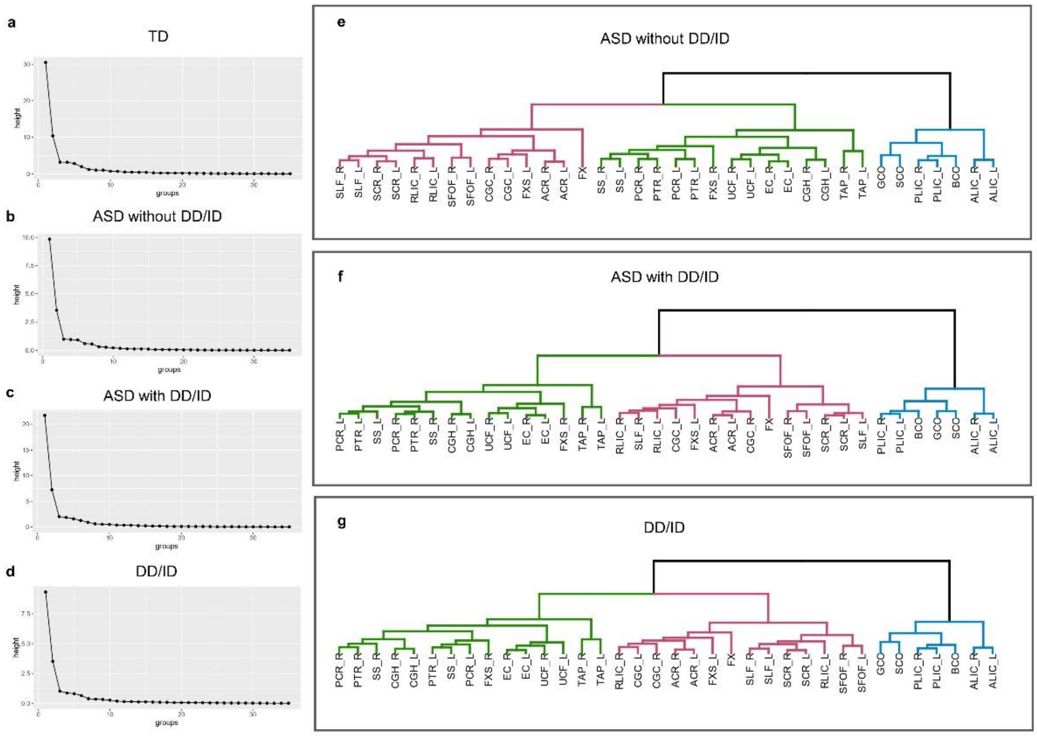

**Supplementary Fig. 4a. Clustering results and developmental curves of each validation group in Splitting analyses in ASD with DD/ID and TD.** ASD, autism spectrum disorder. DD/ID, developmental delay/intellectual disability. TD, typical developing.

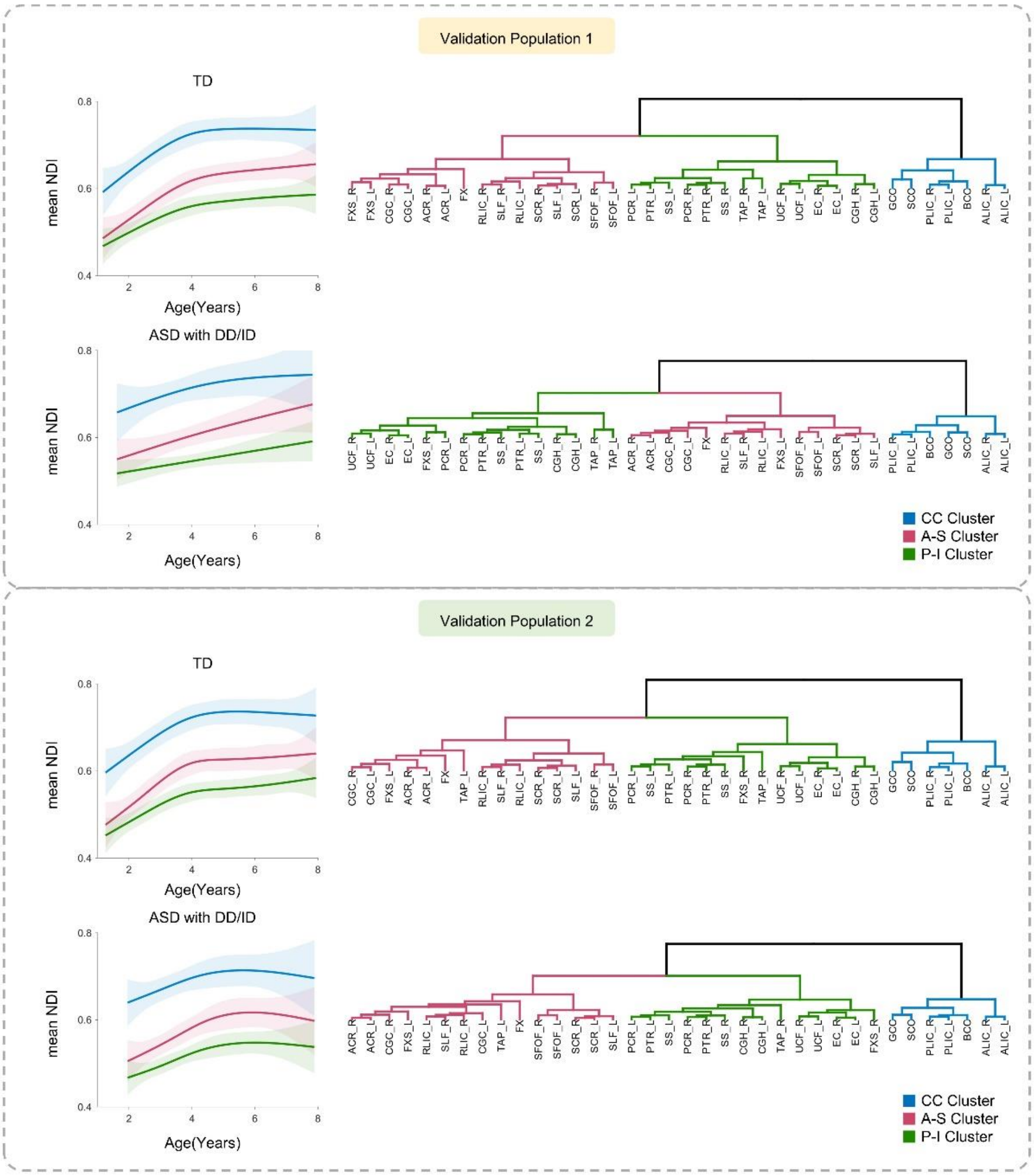

**Supplementary Fig. 4b. Growth rates and developmental stages of each validation group in Splitting analyses in ASD with DD/ID and TD.** ASD, autism spectrum disorder. DD/ID, developmental delay/intellectual disability. TD, typical developing.

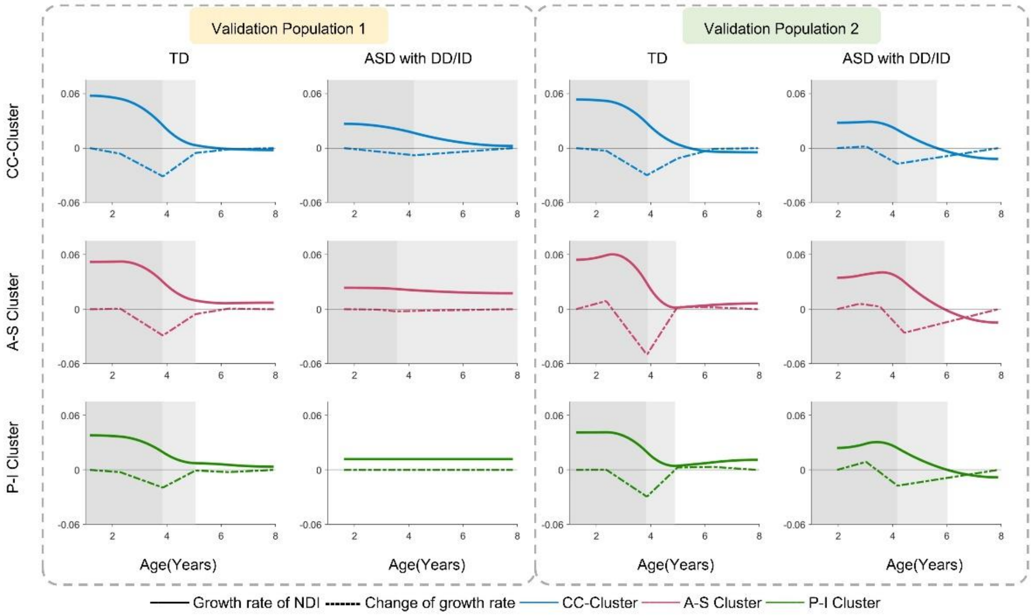

**Supplementary Fig. 5a. Clustering results and developmental curves in male-only sample of all groups.** ASD, autism spectrum disorder. DD/ID, developmental delay/intellectual disability. TD, typical developing.

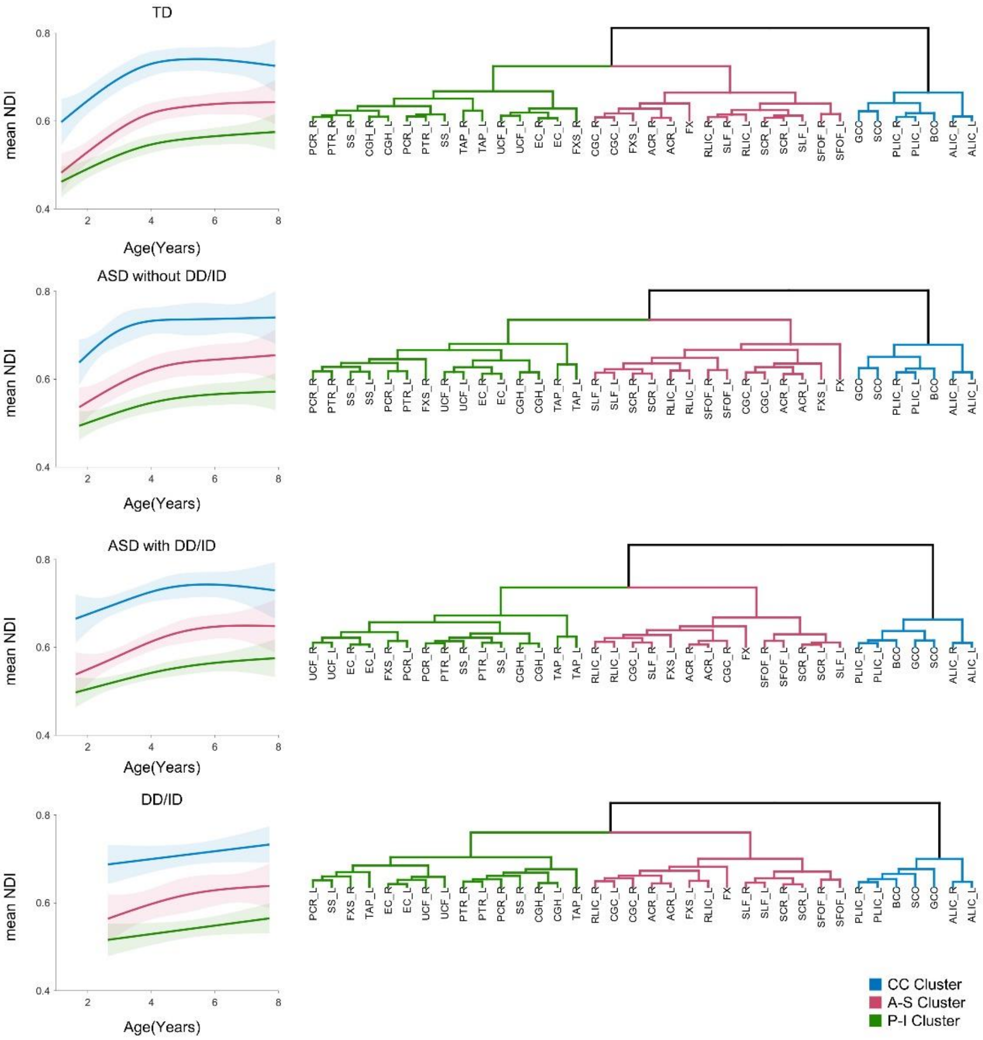

**Supplementary Fig. 5b. Growth rates and developmental stages in male-only sample of all groups.** ASD, autism spectrum disorder. DD/ID, developmental delay/intellectual disability. TD, typical developing.

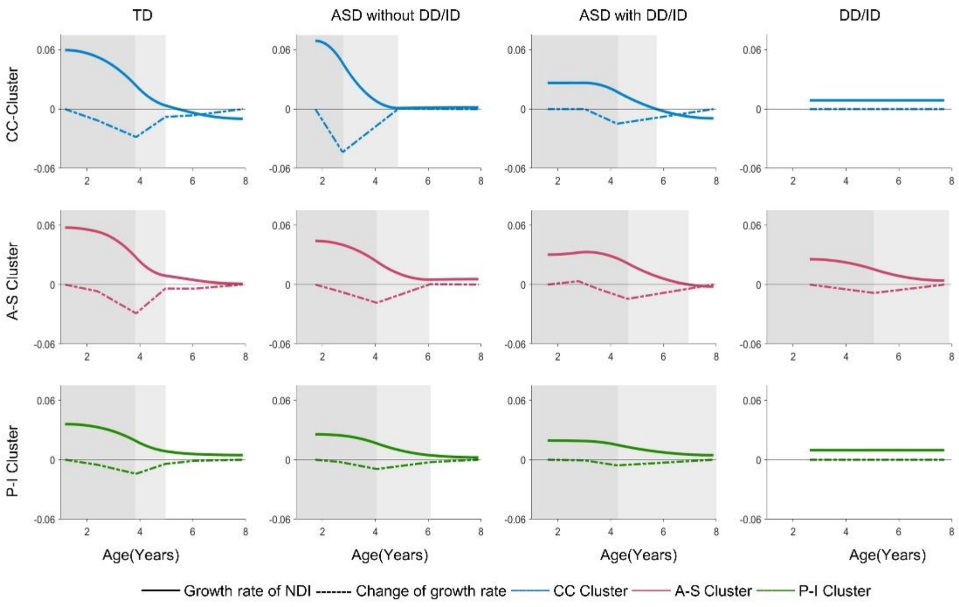

**Supplementary Fig. 6a. Developmental curves of FA in white matter clusters of each group.**

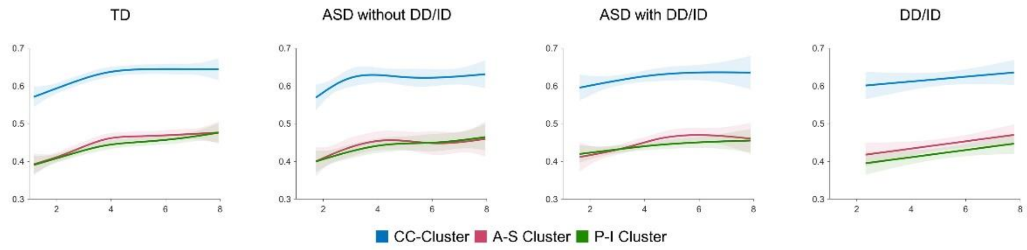


ASD, autism spectrum disorder. DD/ID, developmental delay/intellectual disability. TD, typical developing.

**Supplementary Fig. 6b. Growth rates of early white matter maturation based on FA.**

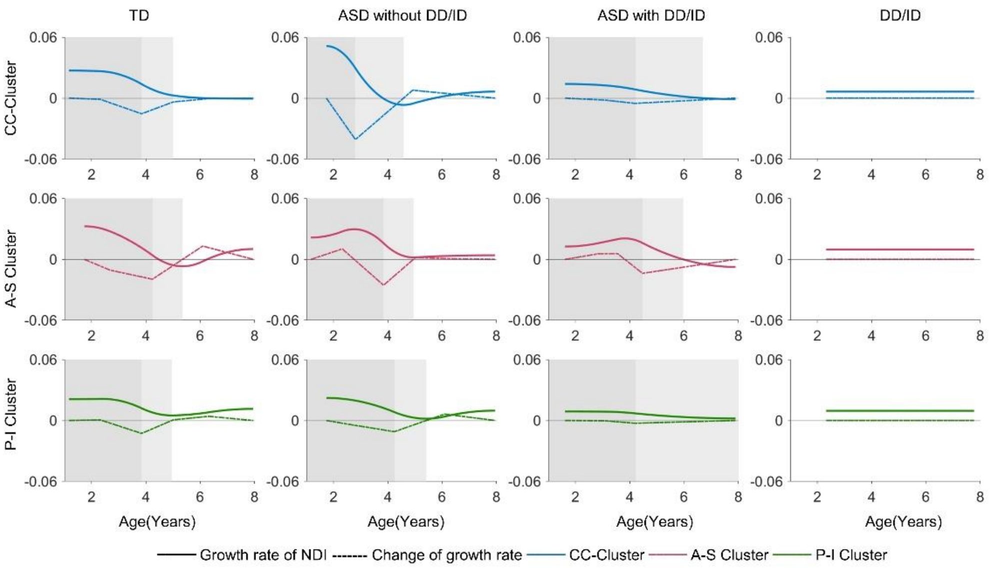


ASD, autism spectrum disorder. DD/ID, developmental delay/intellectual disability. TD, typical developing. The growth rates of FA were calculated by the first derivative of developmental curves (solid line). The changes of rates were represented by the derivative of growth rate (dotted line). According to the changes of rates for all white matter clusters, three developmental stages were defined: fast, moderate and slow stages (shadow areas) except DD/ID group.

### Supplementary Table 1. The summary of the demographics in this study.

| | ASD with DD/ID (A) | ASD without DD/ID (B) | DD/ID (C) | TD (D) | Post hoc[a] |
|---|---|---|---|---|---|
| **Number** | 105 | 51 | 48 | 160 | n.a. |
| **Age (years, mean ± std)** | 3.93±1.51 | 4.32±1.95 | 5.24±1.73 | 4.39±1.90 | C>A,C |
| **Sex (% male)** | 87.62 | 82.35 | 77.08 | 49.38 | A,B,C>D |
| **Absolute RMS (mm, mean ± std)** | 0.43±0.15 | 0.45±0.23 | 0.45±0.28 | 0.56±0.63 | n.s |
| **Relative RMS (mm, mean ± std)** | 0.10±0.03 | 0.10±0.02 | 0.10±0.03 | 0.11±0.05 | n.s |

ASD, autism spectrum disorder. DD/ID, developmental delay/intellectual disability. TD, typical developing. Absolute RMS, root-mean-square displacement of the mean absolute intervolume displacement. Relative RMS, root-mean-square displacement of the mean relative intervolume displacement. [a] Tukey's honest significant difference criterion, $p$<0.05. n.a., not applicable. n.s., not significant.

**Supplementary Table 2. Clinical measurements of ASD with DD/ID, ASD without DD/ID, DD/ID and TD.**

| | ASD with DD/ID (A) | ASD without DD/ID (B) | DD/ID (C) | TD (D) | Post hoc[a] |
|---|---|---|---|---|---|
| **ADOS** | | | | | |
| ADOS_SA CSS | 8.01 ± 2.06 | 7.56 ± 1.63 | 3.42 ± 1.47 | 1.68 ± 1.08 | A,B>C>D |
| ADOS_RRB CSS | 5.73 ± 1.99 | 4.42 ± 2.55 | 1.65 ± 1.41 | 1.15 ± 0.71 | A>B>C,D |
| **SRS** | | | | | |
| Social Awareness | 11.44 ± 2.90 | 10.33 ± 2.72 | 10.33 ± 3.60 | 6.71 ± 2.96 | A,B,C>D |
| Social Cognition | 17.42 ± 4.74 | 14.92 ± 4.66 | 15.43 ± 6.00 | 8.09 ± 4.80 | A>B<br>A,B,C>D |
| Social Communication | 32.07 ± 8.62 | 26.14 ± 9.40 | 23.20 ± 8.90 | 10.24 ± 7.65 | A>B,C>D |
| Social Motivation | 15.30 ± 4.91 | 12.76 ± 4.50 | 11.58 ± 5.48 | 7.18 ± 3.36 | |
| Autistic Mannerisms | 13.15 ± 5.98 | 10.69 ± 5.96 | 9.23 ± 6.31 | 3.59 ± 3.33 | |
| Total | 89.44 ± 21.90 | 74.90 ± 22.72 | 69.75 ± 27.15 | 35.53 ± 18.93 | |
| **CARS** | | | | | |
| CARS_SUM | 37.14 ± 3.87 | 33.28 ± 4.09 | 25.30 ± 3.10 | 17.51 ± 3.82 | A>B>C>D |
| Human Relatedness | 2.77 ± 0.38 | 2.42 ± 0.41 | 1.80 ± 0.38 | 1.24 ± 0.43 | |
| Imitation | 2.39 ± 0.52 | 2.02 ± 0.59 | 1.65 ± 0.42 | 1.14 ± 0.34 | |
| Affect | 2.76 ± 0.36 | 2.49 ± 0.38 | 1.88 ± 0.43 | 1.23 ± 0.39 | |
| Use Of Body | 2.52 ± 0.43 | 2.16 ± 0.45 | 1.86 ± 0.46 | 1.20 ± 0.39 | |
| Relation To Objects | 2.71 ± 0.38 | 2.26 ± 0.43 | 1.90 ± 0.36 | 1.17 ± 0.38 | |
| Adaptation To Change | 2.37 ± 0.48 | 2.29 ± 0.53 | 1.59 ± 0.49 | 1.18 ± 0.37 | A,B>C>D |
| Visual Responsiveness | 2.57 ± 0.50 | 2.34 ± 0.43 | 1.54 ± 0.45 | 1.11 ± 0.30 | A>B>C>D |
| Auditory Responsiveness | 2.46 ± 0.44 | 2.16 ± 0.57 | 1.41 ± 0.43 | 1.10 ± 0.29 | |
| Near Receptor Responsiveness | 2.40 ± 0.45 | 2.16 ± 0.42 | 1.56 ± 0.52 | 1.23 ± 0.42 | |
| Anxiety Reaction | 1.87 ± 0.44 | 1.66 ± 0.47 | 1.15 ± 0.31 | 1.07 ± 0.23 | |
| Verbal Communication | 2.81 ± 0.31 | 2.50 ± 0.42 | 2.34 ± 0.51 | 1.25 ± 0.49 | A>B,C>D |
| Nonverbal Communication | 2.34 ± 0.56 | 1.88 ± 0.49 | 1.37 ± 0.45 | 1.06 ± 0.22 | A>B>C>D |
| Activity Level | 2.26 ± 0.48 | 2.28 ± 0.44 | 1.82 ± 0.51 | 1.22 ± 0.44 | A,B>C>D |
| Intellectual Consistency | 2.25 ± 0.37 | 2.42 ± 0.51 | 2.15± 0.36 | 1.27 ± 0.54 | A,B,C>D<br>B>C |
| Global Impression | 2.71 ± 0.42 | 2.29 ± 0.49 | 1.29± 0.47 | 1.09 ± 0.27 | A>B>C>D |
| **GDS - DQ** | (n=68) | (n=24) | (n=13) | (n=50) | |
| Gross Motor DQ | 81.22 ± 13.05 | 97.00 ± 8.62 | 76.08 ± 13.25 | 99.23 ± 14.13 | B,D>A,C |
| Fine Motor DQ | 66.77 ± 15.63 | 95.40 ± 8.88 | 59.75 ± 21.41 | 102.34 ± 12.78 | D>A,B,C<br>B>A,C |
| Adaption DQ | 62.31 ± 14.35 | 87.67 ± 10.79 | 59.46 ± 11.13 | 101.89 ± 13.05 | |
| Language DQ | 42.86 ± 13.40 | 62.46 ± 15.40 | 41.15 ± 8.67 | 95.06 ± 20.90 | |
| Personal-Social DQ | 57.66 ± 9.94 | 77.08 ± 9.40 | 56.77 ± 8.72 | 96.06 ± 15.82 | |
| Average DQ | 61.68 ± 8.40 | 83.04± 5.99 | 58.75 ± 8.00 | 99.02 ± 12.96 | |
| **WPPSI/WISC-R - IQ** | (n=33) | (n=25) | (n=34) | (n=86) | |
| Verbal IQ | 52.45 ± 9.18 | 85.52 ± 14.96 | 56.00 ± 8.75 | 105.44 ± 18.13 | D>A,B,C<br>B>A,C |
| Performance IQ | 59.30 ± 12.50 | 96.28 ± 18.22 | 60.94 ± 11.17 | 110.08 ± 15.43 | |
| Full IQ | 51.79 ± 8.44 | 89.60 ± 12.13 | 53.56 ± 7.50 | 108.45 ± 16.92 | |

ASD, autism spectrum disorder. DD/ID, developmental delay/intellectual disability. TD, typical developing. ADOS, Autism Diagnostic Observation Schedule. SRS, Social Responsiveness Scale. CARS, Childhood Autism Rating Scale. WPPSI, Wechsler Preschool and Primary Scale of Intelligence. WISC-R, Wechsler Intelligence Scale for Children-Revised. DQ, developmental quotient. IQ, intelligence quotient. [a] Tukey's honest significant difference criterion, $p<0.05$.

**Supplementary Table 3. The list of cerebral white matter regions in each white matter cluster.**

| | | White matter regions | |
|---|---|---|---|
| **CC Cluster** | Corpus Callosum Cluster | GCC | Genu of corpus callosum |
| | | BCC | Body of corpus callosum |
| | | SCC | Splenium of corpus callosum |
| | | ALIC R&L | Right and left anterior limb of internal capsule |
| | | PLIC R&L | Right and left posterior limb of internal capsule |
| **A-S Cluster** | Anterior-Superior Cluster | ACR R&L | Right and left anterior corona radiata |
| | | SCR R&L | Right and left superior corona radiata |
| | | CGC R&L | Right and left cingulum (cingulate gyrus) |
| | | SLF R&L | Right and left superior longitudinal fasciculus |
| | | SFOF R&L | Right and left superior fronto-occipital fasciculus |
| | | RLIC R&L | Right and left retrolenticular part of internal capsule |
| | | FXS L | Left fornix (cres) / Stria terminalis |
| | | FX | Fornix |
| **P-I Cluster** | Posterior-Inferior Cluster | PCR R&L | Right and left posterior corona radiata |
| | | PTR R&L | Right and left posterior thalamic radiation |
| | | SS R&L | Right and left sagittal stratum |
| | | CGH R&L | Right and left cingulum (cingulate gyrus) |
| | | UCF R&L | Right and left uncinate fasciculus |
| | | EC R&L | Right and left external capsule |
| | | TAP R&L | Right and left tapetum |
| | | FXS R | Right fornix (cres) / Stria terminalis |

**Supplementary Table 4. Two special timepoints used to define the developmental stages in each white matter cluster of all four groups.**

| | | TD | ASD without DD/ID | ASD with DD/ID | DD/ID |
|---|---|---|---|---|---|
| CC Cluster | Point1 (Years) | 3.84 | 2.80 | 4.21 | n.a. |
| | Point2 (Years) | 5.00 | 4.71 | 6.07 | n.a. |
| A-S Cluster | Point1 (Years) | 3.84 | 4.24 | 4.21 | n.a. |
| | Point2 (Years) | 4.76 | 5.60 | 7.89 | n.a. |
| P-I Cluster | Point1 (Years) | 3.84 | 4.24 | 4.21 | n.a. |
| | Point2 (Years) | 4.72 | 6.13 | 7.89 | n.a. |

n.a., not applicable.

**Supplementary Table 5. Group comparisons of growth rates of white matter clusters at each stage and across the entire age range.**

| | | CC Cluster | | A-S Cluster | | P-I Cluster | |
|---|---|---|---|---|---|---|---|
| | | β | *p* | β | *p* | β | *p* |
| **The fast stage** | ASD without DD/ID-TD | 0.013371 | **5.9E-28[a]** | -0.00817 | **2.05E-07[a]** | -0.01173 | **2.23E-33[a]** |
| | ASD with DD/ID-TD | -0.01902 | **1.44E-59[a]** | -0.01808 | **3.29E-32[a]** | -0.01698 | **6.19E-63[a]** |
| | ASD with DD/ID-ASD without DD/ID | -0.03239 | **2.37E-67[a]** | -0.0099 | **4.79E-10[a]** | -0.00525 | **7.71E-11[a]** |
| **The moderate-slow stage** | ASD without DD/ID-TD | 0.001831 | 0.110961 | 0.004671 | **2.85E-05[a]** | 0.001906 | **0.003639[a]** |
| | ASD with DD/ID-TD | 0.001239 | 0.272652 | 0.0056 | **3E-08[a]** | 0.003477 | **8.97E-09[a]** |
| | ASD with DD/ID- ASD without DD/ID | -0.00059 | 0.669282 | 0.000928 | 0.465322 | 0.001571 | **0.039301[a]** |
| **The entire age range** | ASD without DD/ID-TD | 0.004264 | **0.018743[a]** | 0.001585 | 0.352232 | -0.00265 | **0.01951a** |
| | ASD with DD/ID-TD | -0.00773 | **9.58E-08[a]** | -0.00339 | **0.011666[a]** | -0.00563 | **7.27E-10[a]** |
| | DD/ID-TD | -0.00366 | 0.050808 | -0.00327 | 0.063838 | -0.00505 | **2.13E-05[a]** |
| | ASD with DD/ID-ASD without DD/ID | -0.01199 | **1.23E-09[a]** | -0.00497 | **0.006308[a]** | -0.00297 | **0.014001[a]** |
| | DD/ID-ASD without DD/ID | -0.00793 | **0.000565[a]** | -0.00486 | **0.0242[a]** | -0.00239 | 0.09483 |
| | ASD with DD/ID-DD/ID | -0.00406 | 0.043429 | -0.00011 | 0.952008 | -0.00058 | 0.64376 |

The moderate and slow developmental stages were combined as the moderate-slow stage. [a] Fasle discovery rate (FDR) correction with Benjamini-Hochberg procedure for multiple comparisons, $p$>0.05.

**Supplementary Table 6. Comparisons of mean NDI in white matter clusters between fast stage and moderate-slow stage for ASD subgroups and TD group.**

| | CC Cluster | | A-S Cluster | | P-I Cluster | |
|---|---|---|---|---|---|---|
| | **β** | ***p*** | **β** | ***p*** | **β** | ***p*** |
| **ASD without DD/ID** | 0.037016 | **0.033264** | 0.004874 | 0.797557 | 0.006428 | 0.630792 |
| **ASD with DD/ID** | -0.00311 | 0.841027 | -0.00075 | 0.95576 | 0.001141 | 0.913201 |
| **TD** | 0.044741 | **0.000504** | 0.043795 | **0.000113** | 0.024447 | **0.006569** |

The moderate and slow developmental stages were combined as the moderate-slow stage. [a] Fasle discovery rate (FDR) correction with Benjamini-Hochberg procedure for multiple comparisons, $p$>0.05.

**Supplementary Table 7. Group comparisons of mean NDI in white matter clusters at each stage and across entire age range.**

| | | CC Cluster | | A-S Cluster | | P-I Cluster | |
|---|---|---|---|---|---|---|---|
| | | **β** | ***p*** | **β** | ***p*** | **β** | ***p*** |
| **The fast stage** | ASD without DD/ID v.s. TD | 0.00972169 | 0.341139 | 0.006671 | 0.39742 | 0.002484 | 0.695992 |
| | ASD with DD/ID v.s. TD | 0.00898923 | 0.2456229 | 0.004804 | 0.455921 | 0.002233 | 0.667381 |
| | ASD with DD/ID v.s. ASD without DD/ID | -0.0007325 | 0.9439742 | -0.00187 | 0.798973 | -0.00025 | 0.966231 |
| **The moderate-slow stage** | ASD without DD/ID v.s. TD | -0.01379 | 0.071415 | -0.00555 | 0.46457 | -0.00585 | 0.348602 |
| | ASD with DD/ID v.s. TD | -0.00038 | 0.960276 | 0.001719 | 0.806759 | -0.00244 | 0.672649 |
| | ASD with DD/ID v.s. ASD without DD/ID | 0.013406 | 0.131341 | 0.00727 | 0.404186 | 0.003406 | 0.633898 |
| **The entire age range** | ASD without DD/ID v.s. TD | -0.0043 | 0.529346 | -3.9E-06 | 0.9995 | -0.00213 | 0.658298 |
| | ASD with DD/ID v.s. TD | 0.011951 | 0.032302 | 0.008525 | 0.094227 | 0.003514 | 0.369966 |
| | DD/ID v.s. TD | -0.01432 | 0.040434 | -0.01225 | 0.054773 | -0.01208 | **0.014155**[a] |
| | ASD with DD/ID v.s. ASD without DD/ID | 0.016254 | 0.021106 | 0.008529 | 0.184217 | 0.005644 | 0.254019 |
| | DD/ID v.s. ASD without DD/ID | -0.01001 | 0.229888 | -0.01225 | 0.108183 | -0.00995 | 0.090418 |
| | ASD with DD/ID v.s. DD/ID | 0.026267 | **0.000384**[a] | 0.020777 | **0.002062**[a] | 0.015595 | **0.002682**[a] |

The moderate and slow developmental stages were combined as the moderate-slow stage. [a] Fasle discovery rate (FDR) correction with Benjamini-Hochberg procedure for multiple comparisons, $p$>0.05.

**Supplementary Table 8a. Correlations between NDI of white matter clusters and social symptoms and cognitive performances of ASD with DD/ID during the fast stage.**

| | CC Cluster | | A-S Cluster | | P-I Cluster | |
|---|---|---|---|---|---|---|
| | **r** | ***p*** | **r** | ***p*** | **r** | ***p*** |
| CARS Sum | -0.01144 | 0.927926 | 0.108917 | 0.387781 | -0.04646 | 0.71325 |
| CARS Human Relatedness | 0.07686 | 0.54283 | 0.186803 | 0.136228 | 0.09213 | 0.465439 |
| CARS Imitation | 0.045134 | 0.721091 | 0.16454 | 0.190273 | 0.100506 | 0.425683 |
| CARS Affect | 0.048967 | 0.698494 | 0.095548 | 0.448979 | -0.00382 | 0.975938 |
| CARS Use Of Body | 0.084734 | 0.502154 | 0.194712 | 0.120118 | 0.049378 | 0.696085 |
| CARS Relation To Objects | 0.266373 | 0.031968 | 0.296667 | 0.016409 | 0.237051 | 0.057265 |
| CARS Adaptation To Change | -0.04861 | 0.700572 | -0.03837 | 0.761568 | -0.06785 | 0.591247 |
| CARS Visual Responsiveness | 0.002153 | 0.986422 | 0.116438 | 0.355648 | -0.01249 | 0.921346 |
| CARS Auditory Responsiveness | -0.0373 | 0.768025 | 0.090797 | 0.471948 | -0.08736 | 0.488949 |
| CARS Near Receptor Responsiveness | -0.1798 | 0.151807 | -0.05516 | 0.662537 | -0.12289 | 0.329455 |
| CARS Anxiety Reaction | 0.021013 | 0.868042 | 0.05314 | 0.674185 | -0.07692 | 0.54253 |
| CARS Verbal Communication | -0.03774 | 0.765335 | 0.022125 | 0.861129 | 0.02671 | 0.832727 |
| CARS Nonverbal Communication | -0.00272 | 0.98285 | 0.122528 | 0.330874 | 0.04609 | 0.715433 |
| CARS Activity Level | -0.07132 | 0.572396 | 0.022075 | 0.861437 | -0.07125 | 0.572742 |
| CARS Intellectual Consistency | 0.053753 | 0.675646 | 0.127948 | 0.317633 | 0.119347 | 0.351516 |
| CARS Global Impression | 0.00119 | 0.992496 | 0.121256 | 0.335955 | 0.017499 | 0.889963 |
| ADOS_SA_CSS | 0.034999 | 0.778578 | 0.027953 | 0.822335 | -0.00546 | 0.965027 |
| ADOS_RRB_CSS | -0.14883 | 0.260588 | -0.15253 | 0.2488 | -0.16623 | 0.208295 |
| Gross Motor DQ | -0.07149 | 0.574546 | -0.1108 | 0.383398 | -0.14535 | 0.251801 |
| Fine Motor DQ | -0.25306 | 0.142445 | **-0.44326** | **0.007656**[a] | -0.365 | 0.031086 |
| Adaptive DQ | -0.12852 | 0.311504 | -0.25157 | 0.044931 | **-0.34292** | **0.005538**[a] |
| Language DQ | -0.12302 | 0.332811 | -0.15319 | 0.226852 | -0.09268 | 0.466352 |
| Personal-Social DQ | -0.07973 | 0.534474 | -0.08364 | 0.514604 | -0.11309 | 0.377521 |
| Average DQ | -0.19145 | 0.129652 | -0.2976 | 0.01693 | **-0.31344** | **0.011668**[a] |

The moderate and slow developmental stages were combined as the moderate-slow stage. [a] Fasle discovery rate (FDR) correction with Benjamini-Hochberg procedure for multiple comparisons, *p*>0.05.

**Supplementary Table 8b. Correlations between NDI of white matter clusters and social symptoms and cognitive performances of ASD with DD/ID during the moderate-slow stage.**

| | CC Cluster | | A-S Cluster | | P-I Cluster | |
|---|---|---|---|---|---|---|
| | r | *p* | r | *p* | r | *p* |
| CARS Sum | -0.30811 | 0.097627 | -0.2923 | 0.117017 | -0.25532 | 0.173285 |
| CARS Human Relatedness | -0.3293 | 0.075578 | -0.33459 | 0.070721 | -0.35272 | 0.055903 |
| CARS Imitation | -0.0014 | 0.994146 | -0.126 | 0.507028 | -0.02098 | 0.912393 |
| CARS Affect | -0.08053 | 0.672267 | -0.1025 | 0.589886 | -0.03488 | 0.854805 |
| CARS Use Of Body | -0.10097 | 0.595488 | 0.124306 | 0.512811 | 0.116877 | 0.538506 |
| CARS Relation To Objects | -0.24829 | 0.185846 | -0.08615 | 0.650798 | -0.01726 | 0.927859 |
| CARS Adaptation To Change | -0.09439 | 0.61979 | -0.13059 | 0.491553 | -0.11452 | 0.54679 |
| CARS Visual Responsiveness | -0.12513 | 0.517776 | -0.23768 | 0.214418 | -0.18154 | 0.345942 |
| CARS Auditory Responsiveness | -0.15229 | 0.430328 | -0.065 | 0.737623 | -0.00022 | 0.99908 |
| CARS Near Receptor Responsiveness | -0.30309 | 0.109991 | -0.07963 | 0.681341 | -0.22749 | 0.235291 |
| CARS Anxiety Reaction | -0.21122 | 0.27139 | -0.06768 | 0.727203 | -0.13679 | 0.479216 |
| CARS Verbal Communication | -0.37975 | 0.042167 | -0.212 | 0.26957 | -0.22967 | 0.230722 |
| CARS Nonverbal Communication | -0.28278 | 0.137184 | -0.33793 | 0.072983 | -0.24656 | 0.197269 |
| CARS Activity Level | -0.24963 | 0.191564 | -0.05714 | 0.76843 | -0.12469 | 0.519281 |
| CARS Intellectual Consistency | 0.272963 | 0.159907 | -0.06394 | 0.746524 | 0.130074 | 0.509434 |
| CARS Global Impression | -0.20844 | 0.277894 | -0.26042 | 0.172449 | -0.12971 | 0.50245 |
| ADOS_SA_CSS | -0.1999 | 0.29849 | -0.20207 | 0.293158 | -0.21833 | 0.255202 |
| ADOS_RRB_CSS | -0.11557 | 0.550523 | -0.05496 | 0.777033 | 0.038302 | 0.843623 |
| Verbal IQ | 0.321566 | 0.10192 | 0.379859 | 0.050657 | 0.355578 | 0.068725 |
| Performance IQ | 0.204194 | 0.306955 | 0.300344 | 0.127964 | 0.156585 | 0.435409 |
| Full IQ | 0.326711 | 0.09625 | 0.3511 | 0.072542 | 0.318771 | 0.105102 |
| SRS Total | -0.00697 | 0.970842 | 0.168889 | 0.372302 | 0.082227 | 0.665761 |
| SRS Social Awareness | -0.09765 | 0.607716 | 0.202682 | 0.28275 | 0.021678 | 0.909474 |
| SRS Social Cognition | 0.132666 | 0.484636 | 0.277928 | 0.136997 | 0.251827 | 0.179447 |
| SRS Social Communication | -0.07606 | 0.689528 | -0.05859 | 0.758423 | -0.083 | 0.662792 |
| SRS Social Motivation | -0.0082 | 0.965681 | 0.048109 | 0.800689 | 0.018566 | 0.922426 |
| SRS Autistic Mannerisms | 0.03958 | 0.835496 | 0.301013 | 0.106007 | 0.183085 | 0.33284 |

The moderate and slow developmental stages were combined as the moderate-slow stage. [a] Fasle discovery rate (FDR) correction with Benjamini-Hochberg procedure for multiple comparisons, $p>0.05$.

**Supplementary Table 8c. Correlations between NDI of white matter clusters and social symptoms and cognitive performances of ASD without DD/ID at the fast stage.**

| | CC Cluster | | A-S Cluster | | P-I Cluster | |
|---|---|---|---|---|---|---|
| | **r** | ***p*** | **r** | ***p*** | **r** | ***p*** |
| CARS Sum | -0.39239 | 0.184779 | -0.23257 | 0.310331 | 0.074039 | 0.749762 |
| CARS Human Relatedness | -0.00363 | 0.990605 | -0.19626 | 0.393856 | -0.01593 | 0.945345 |
| CARS Imitation | -0.2911 | 0.334569 | -0.19718 | 0.3916 | 0.182717 | 0.427923 |
| CARS Affect | -0.2676 | 0.376749 | -0.06108 | 0.792551 | 0.258085 | 0.258669 |
| CARS Use Of Body | -0.71105 | 0.006432 | -0.25068 | 0.27307 | -0.31926 | 0.158341 |
| CARS Relation To Objects | -0.34883 | 0.242739 | -0.06843 | 0.768205 | 0.110304 | 0.634088 |
| CARS Adaptation To Change | 0.129774 | 0.672621 | 0.118821 | 0.607961 | 0.09115 | 0.69436 |
| CARS Visual Responsiveness | -0.04137 | 0.893243 | -0.18542 | 0.420997 | 0.101489 | 0.661578 |
| CARS Auditory Responsiveness | -0.12931 | 0.673726 | -0.06841 | 0.768267 | -0.05691 | 0.806437 |
| CARS Near Receptor Responsiveness | -0.39818 | 0.177814 | 0.051041 | 0.82609 | 0.067758 | 0.770419 |
| CARS Anxiety Reaction | 0.035284 | 0.908894 | -0.18829 | 0.413723 | -0.04344 | 0.851691 |
| CARS Verbal Communication | -0.24279 | 0.424151 | -0.2663 | 0.24328 | -0.14696 | 0.524974 |
| CARS Nonverbal Communication | -0.22518 | 0.459496 | -0.1562 | 0.498961 | -0.11304 | 0.625653 |
| CARS Activity Level | -0.06465 | 0.833804 | 0.100068 | 0.666049 | 0.175658 | 0.446282 |
| CARS Intellectual Consistency | 0.39988 | 0.175793 | 0.157807 | 0.494492 | 0.273381 | 0.230494 |
| CARS Global Impression | -0.5259 | 0.064895 | -0.35759 | 0.111504 | 0.041943 | 0.856746 |
| ADOS_SA_CSS | -0.14299 | 0.641202 | -0.211 | 0.358565 | 0.000293 | 0.998996 |
| ADOS_RRB_CSS | -0.38327 | 0.196108 | -0.26971 | 0.237058 | -0.10099 | 0.66316 |
| Gross Motor DQ | 0.451978 | 0.104694 | 0.459009 | 0.048059 | 0.456201 | 0.049621 |
| Fine Motor DQ | 0.339764 | 0.306648 | 0.307422 | 0.357767 | 0.240566 | 0.476121 |
| Adaptive DQ | 0.379356 | 0.180979 | 0.191481 | 0.432284 | 0.198951 | 0.41419 |
| Language DQ | -0.05172 | 0.860618 | -0.23546 | 0.331844 | -0.31291 | 0.192095 |
| Personal-Social DQ | -0.00196 | 0.994686 | -0.15723 | 0.52033 | -0.05175 | 0.833355 |
| Average DQ | 0.295943 | 0.304267 | -0.01359 | 0.955965 | 0.005914 | 0.980829 |

The moderate and slow developmental stages were combined as the moderate-slow stage. [a] Fasle discovery rate (FDR) correction with Benjamini-Hochberg procedure for multiple comparisons, $p$>0.05.

**Supplementary Table 8d. Correlations between NDI of white matter clusters and social symptoms and cognitive performances of ASD without DD/ID during the moderate-slow stage.**

| | **CC Cluster** | | **A-S Cluster** | | **P-I Cluster** | |
|---|---|---|---|---|---|---|
| | **r** | ***p*** | **r** | ***p*** | **r** | ***p*** |
| CARS Sum | 0.123744 | 0.522476 | 0.117084 | 0.613253 | 0.236143 | 0.302754 |
| CARS Human Relatedness | 0.217323 | 0.257449 | 0.272756 | 0.231604 | 0.395822 | 0.075703 |
| CARS Imitation | -0.04439 | 0.819146 | 0.080938 | 0.727265 | 0.147515 | 0.523401 |
| CARS Affect | -0.07752 | 0.689394 | -0.31554 | 0.163514 | -0.12748 | 0.581857 |
| CARS Use Of Body | 0.015509 | 0.936357 | 0.034984 | 0.880335 | 0.056969 | 0.806243 |
| CARS Relation To Objects | -0.12405 | 0.521437 | 0.020246 | 0.930588 | 0.095433 | 0.680711 |
| CARS Adaptation To Change | -0.16766 | 0.384657 | -0.01447 | 0.950362 | 0.11483 | 0.620149 |
| CARS Visual Responsiveness | 0.12971 | 0.502458 | 0.285677 | 0.209357 | 0.287554 | 0.206249 |
| CARS Auditory Responsiveness | 0.216471 | 0.259365 | -0.08382 | 0.717928 | 0.076572 | 0.741478 |
| CARS Near Receptor Responsiveness | -0.14593 | 0.450043 | -0.27937 | 0.220035 | -0.12606 | 0.586116 |
| CARS Anxiety Reaction | -0.12277 | 0.525779 | 0.584998 | 0.005343 | 0.563182 | 0.007853 |
| CARS Verbal Communication | 0.168148 | 0.383259 | 0.22184 | 0.333813 | 0.278486 | 0.221555 |
| CARS Nonverbal Communication | 0.030868 | 0.873704 | -0.06081 | 0.793445 | 0.19401 | 0.399411 |
| CARS Activity Level | -0.13369 | 0.489329 | 0.21621 | 0.346537 | 0.070688 | 0.760763 |
| CARS Intellectual Consistency | 0.352134 | 0.066102 | 0.54913 | 0.012153 | 0.341323 | 0.140802 |
| CARS Global Impression | -0.02574 | 0.894557 | 0.042247 | 0.855719 | 0.194798 | 0.39746 |
| ADOS_SA_CSS | -0.06132 | 0.751997 | 0.164869 | 0.475121 | 0.213452 | 0.352871 |
| ADOS_RRB_CSS | -0.29277 | 0.123249 | -0.01389 | 0.952365 | -0.28761 | 0.20616 |
| Verbal IQ | -0.50964 | 0.018275 | -0.32861 | 0.183056 | -0.53279 | 0.022814 |
| Performance IQ | 0.324774 | 0.150874 | 0.514375 | 0.028963 | 0.371603 | 0.12892 |
| Full IQ | -0.13083 | 0.571907 | 0.150832 | 0.550227 | -0.12605 | 0.618223 |
| SRS Total | -0.23185 | 0.299174 | -0.12526 | 0.609375 | -0.2297 | 0.344146 |
| SRS Social Awareness | **-0.5652** | **0.006124**[a] | -0.473 | 0.040826 | -0.47918 | 0.037909 |
| SRS Social Cognition | -0.12994 | 0.564375 | -0.10339 | 0.673619 | -0.00913 | 0.970412 |
| SRS Social Communication | -0.17502 | 0.435954 | 0.010011 | 0.967556 | -0.13158 | 0.591305 |
| SRS Social Motivation | 0.091237 | 0.686358 | 0.082519 | 0.736986 | -0.11594 | 0.636467 |
| SRS Autistic Mannerisms | -0.26721 | 0.229294 | -0.18176 | 0.456442 | -0.28515 | 0.236688 |

The moderate and slow developmental stages were combined as the moderate-slow stage. [a] Fasle discovery rate (FDR) correction with Benjamini-Hochberg procedure for multiple comparisons, $p>0.05$.

**Supplementary Table 9a. Correlations between NDI of white matter clusters and social symptoms and cognitive performances of ASD without DD/ID across the entire age.**

| | CC Cluster | | A-S Cluster | | P-I Cluster | |
|---|---|---|---|---|---|---|
| | r | *p* | r | *p* | r | *p* |
| CARS Sum | -0.06234 | 0.68065 | -0.00872 | 0.954155 | 0.170067 | 0.258491 |
| CARS Human Relatedness | -0.0121 | 0.936369 | 0.027664 | 0.855191 | 0.107243 | 0.478081 |
| CARS Imitation | -0.06107 | 0.686815 | 0.07801 | 0.606334 | 0.215696 | 0.14996 |
| CARS Affect | -0.19586 | 0.192067 | -0.16965 | 0.259693 | 0.051471 | 0.734075 |
| CARS Use Of Body | -0.18988 | 0.206252 | -0.05822 | 0.70072 | -0.07173 | 0.635691 |
| CARS Relation To Objects | -0.11801 | 0.434743 | -0.05521 | 0.715564 | 0.070218 | 0.642857 |
| CARS Adaptation To Change | -0.10076 | 0.505212 | -0.11265 | 0.456043 | 0.012642 | 0.933544 |
| CARS Visual Responsiveness | 0.114324 | 0.449331 | 0.109533 | 0.468679 | 0.214351 | 0.152581 |
| CARS Auditory Responsiveness | 0.028775 | 0.849443 | -0.09993 | 0.508746 | -0.09888 | 0.513263 |
| CARS Near Receptor Responsiveness | -0.16255 | 0.280438 | -0.09053 | 0.549623 | 0.03938 | 0.794987 |
| CARS Anxiety Reaction | -0.13756 | 0.361955 | 0.067532 | 0.655649 | 0.168376 | 0.263326 |
| CARS Verbal Communication | 0.014427 | 0.924189 | -0.01069 | 0.94377 | 0.082956 | 0.583627 |
| CARS Nonverbal Communication | -0.23667 | 0.113287 | -0.23363 | 0.118125 | -0.06033 | 0.690412 |
| CARS Activity Level | 0.003951 | 0.979207 | 0.098148 | 0.51639 | 0.107766 | 0.475927 |
| CARS Intellectual Consistency | 0.272204 | 0.07046 | 0.292091 | 0.051539 | 0.201672 | 0.184017 |
| CARS Global Impression | -0.22649 | 0.130121 | -0.16564 | 0.271262 | 0.089128 | 0.555837 |
| ADOS_SA_CSS | -0.15379 | 0.307533 | -0.08927 | 0.555195 | 0.073748 | 0.6262 |
| ADOS_RRB_CSS | **-0.37021** | **0.011327**[a] | -0.21176 | 0.157724 | -0.20144 | 0.179459 |
| Gross Motor DQ | 0.399549 | 0.080927 | 0.456531 | 0.043032 | 0.454227 | 0.044231 |
| Fine Motor DQ | 0.339764 | 0.306648 | 0.307422 | 0.357767 | 0.240566 | 0.476121 |
| Adaptive DQ | 0.321875 | 0.166377 | 0.208055 | 0.378735 | 0.19454 | 0.411143 |
| Language DQ | -0.19438 | 0.411537 | -0.26761 | 0.254007 | -0.30067 | 0.197711 |
| Personal-Social DQ | 0.027903 | 0.90704 | -0.12431 | 0.60155 | -0.05543 | 0.816467 |
| Average DQ | 0.143831 | 0.545196 | -0.01102 | 0.963216 | 0.00558 | 0.981374 |
| Verbal IQ | -0.50964 | 0.018275 | -0.36535 | 0.1034 | -0.54457 | 0.010695 |
| Performance IQ | 0.324774 | 0.150874 | 0.423124 | 0.055985 | 0.268423 | 0.239398 |
| Full IQ | -0.13083 | 0.571907 | 0.050017 | 0.829529 | -0.20603 | 0.370248 |
| SRS Total | -0.23185 | 0.299174 | -0.06123 | 0.786637 | -0.09739 | 0.666339 |
| SRS Social Awareness | **-0.5652** | **0.006124**[a] | -0.37979 | 0.081266 | -0.30731 | 0.164162 |
| SRS Social Cognition | -0.12994 | 0.564375 | -0.06906 | 0.760063 | 0.08807 | 0.696735 |
| SRS Social Communication | -0.17502 | 0.435954 | 0.05782 | 0.798274 | -0.03027 | 0.89361 |
| SRS Social Motivation | 0.091237 | 0.686358 | 0.115912 | 0.607477 | -0.01026 | 0.963856 |
| SRS Autistic Mannerisms | -0.26721 | 0.229294 | -0.12782 | 0.570799 | -0.20115 | 0.369387 |

The moderate and slow developmental stages were combined as the moderate-slow stage. [a] Fasle discovery rate (FDR) correction with Benjamini-Hochberg procedure for multiple comparisons, $p$>0.05.

**Supplementary Table 9b. Correlations between NDI and social symptoms and cognitive performance of ASD with DD/ID across the entire age within each cluster.**

| | CC Cluster | | A-S Cluster | | P-I Cluster | |
|---|---|---|---|---|---|---|
| | **r** | ***p*** | **r** | ***p*** | **r** | ***p*** |
| CARS Sum | -0.06944 | 0.494616 | 0.024163 | 0.812347 | -0.09785 | 0.335257 |
| CARS Human Relatedness | -0.04156 | 0.682951 | 0.008087 | 0.93668 | -0.06714 | 0.509072 |
| CARS Imitation | 0.028235 | 0.781453 | 0.125661 | 0.215214 | 0.059572 | 0.558061 |
| CARS Affect | 0.00837 | 0.934465 | 0.042434 | 0.676651 | -0.02831 | 0.780886 |
| CARS Use Of Body | 0.028943 | 0.776118 | 0.146898 | 0.146796 | 0.040926 | 0.687534 |
| CARS Relation To Objects | 0.098334 | 0.332879 | 0.176486 | 0.080558 | 0.102592 | 0.312263 |
| CARS Adaptation To Change | -0.04571 | 0.653225 | -0.05328 | 0.600442 | -0.08747 | 0.389306 |
| CARS Visual Responsiveness | 0.001112 | 0.99133 | 0.051256 | 0.61621 | -0.06729 | 0.5103 |
| CARS Auditory Responsiveness | -0.01474 | 0.885483 | 0.075371 | 0.460752 | -0.0903 | 0.376558 |
| CARS Near Receptor Responsiveness | -0.23142 | 0.021864 | -0.11887 | 0.243693 | -0.15862 | 0.118755 |
| CARS Anxiety Reaction | -0.05576 | 0.585549 | -0.00415 | 0.967669 | -0.0493 | 0.629752 |
| CARS Verbal Communication | -0.09745 | 0.339781 | -0.02398 | 0.814678 | -0.03573 | 0.726851 |
| CARS Nonverbal Communication | -0.01931 | 0.850346 | 0.062292 | 0.542301 | -0.03521 | 0.730693 |
| CARS Activity Level | -0.08278 | 0.41771 | -0.02146 | 0.83386 | -0.07631 | 0.455162 |
| CARS Intellectual Consistency | 0.050477 | 0.627122 | 0.114575 | 0.268899 | 0.092379 | 0.373259 |
| CARS Global Impression | -0.01403 | 0.890975 | 0.051198 | 0.616611 | -0.04701 | 0.645756 |
| ADOS_SA_CSS | -0.02009 | 0.842759 | -0.02914 | 0.7735 | -0.07215 | 0.47565 |
| ADOS_RRB_CSS | -0.16232 | 0.122128 | -0.09685 | 0.358419 | -0.13607 | 0.195912 |
| Gross Motor DQ | -0.08784 | 0.479645 | -0.1108 | 0.383398 | -0.14535 | 0.251801 |
| Fine Motor DQ | -0.25306 | 0.142445 | **-0.44326** | **0.007656**[a] | -0.365 | 0.031086 |
| Adaptive DQ | -0.1321 | 0.2866 | -0.25157 | 0.044931 | **-0.34292** | **0.005538**[a] |
| Language DQ | -0.1658 | 0.179963 | -0.15319 | 0.226852 | -0.09268 | 0.466352 |
| Personal-Social DQ | -0.103 | 0.410507 | -0.08364 | 0.514604 | -0.11309 | 0.377521 |
| Average DQ | -0.21881 | 0.075247 | -0.2976 | 0.01693 | **-0.31344** | **0.011668**[a] |
| Verbal IQ | 0.323374 | 0.087055 | 0.360169 | 0.054956 | 0.385929 | 0.038663 |
| Performance IQ | 0.200312 | 0.297471 | 0.1458 | 0.450451 | 0.278509 | 0.143475 |
| Full IQ | 0.320301 | 0.090273 | 0.305919 | 0.106545 | 0.333985 | 0.076612 |
| SRS Total | -0.05847 | 0.734803 | 0.016175 | 0.9254 | 0.152188 | 0.375574 |
| SRS Social Awareness | -0.09809 | 0.569251 | -0.01136 | 0.947568 | 0.193235 | 0.258824 |
| SRS Social Cognition | 0.080959 | 0.638797 | 0.165324 | 0.335251 | 0.238078 | 0.162037 |
| SRS Social Communication | -0.10624 | 0.537441 | -0.10097 | 0.557901 | -0.0162 | 0.925303 |
| SRS Social Motivation | -0.05602 | 0.745549 | -0.01393 | 0.935752 | 0.078648 | 0.648431 |
| SRS Autistic Mannerisms | -0.03492 | 0.839781 | 0.090083 | 0.601333 | 0.234852 | 0.167971 |

The moderate and slow developmental stages were combined as the moderate-slow stage. [a] Fasle discovery rate (FDR) correction with Benjamini-Hochberg procedure for multiple comparisons, *p*>0.05.

**Supplementary Table 9c. Correlations between NDI of white matter clusters and social symptoms and cognitive performances of DD/ID across the entire age.**

| | CC Cluster | | A-S Cluster | | P-I Cluster | |
|---|---|---|---|---|---|---|
| | r | *p* | r | *p* | r | *p* |
| CARS Sum | 0.126581 | 0.412915 | 0.262585 | 0.085068 | 0.136254 | 0.377827 |
| CARS Human Relatedness | -0.16424 | 0.286736 | -0.04637 | 0.765008 | -0.01235 | 0.936568 |
| CARS Imitation | 0.241413 | 0.114403 | 0.300669 | 0.047351 | 0.070556 | 0.649029 |
| CARS Affect | 0.186555 | 0.225312 | 0.253521 | 0.096812 | 0.179865 | 0.242685 |
| CARS Use Of Body | -0.03728 | 0.810124 | 0.067339 | 0.664063 | 0.049637 | 0.74899 |
| CARS Relation To Objects | 0.103956 | 0.501879 | 0.180935 | 0.239849 | 0.138552 | 0.369758 |
| CARS Adaptation To Change | 0.082617 | 0.593928 | 0.191297 | 0.213531 | 0.057433 | 0.711154 |
| CARS Visual Responsiveness | 0.099946 | 0.518602 | 0.13411 | 0.385447 | 0.058108 | 0.70791 |
| CARS Auditory Responsiveness | -0.20843 | 0.174549 | -0.09965 | 0.519847 | -0.05883 | 0.704435 |
| CARS Near Receptor Responsiveness | 0.205875 | 0.180011 | 0.290704 | 0.055573 | 0.213248 | 0.164573 |
| CARS Anxiety Reaction | 0.235977 | 0.123049 | 0.118322 | 0.444299 | 0.188611 | 0.220152 |
| CARS Verbal Communication | -0.02211 | 0.886723 | 0.028972 | 0.851906 | 0.02001 | 0.897419 |
| CARS Nonverbal Communication | 0.056952 | 0.716794 | 0.119827 | 0.444052 | 0.060146 | 0.701626 |
| CARS Activity Level | 0.25441 | 0.095607 | 0.11848 | 0.443686 | 0.071618 | 0.644098 |
| CARS Intellectual Consistency | -0.05152 | 0.739791 | 0.016445 | 0.915624 | 0.050285 | 0.74582 |
| CARS Global Impression | 0.241406 | 0.114415 | 0.311203 | 0.03976 | 0.209139 | 0.173055 |
| ADOS_SA_CSS | 0.202371 | 0.187701 | 0.300855 | 0.047207 | 0.317337 | 0.03582 |
| ADOS_RRB_CSS | 0.146787 | 0.341703 | 0.175904 | 0.253388 | 0.107158 | 0.488725 |
| Gross Motor DQ | 0.597944 | 0.089002 | 0.747946 | 0.020477 | 0.500181 | 0.170291 |
| Fine Motor DQ | -0.50637 | 0.493627 | 0.936648 | 0.063352 | 0.601103 | 0.398897 |
| Adaptive DQ | -0.07748 | 0.842955 | 0.066858 | 0.864304 | -0.09494 | 0.808033 |
| Language DQ | -0.51228 | 0.158529 | -0.54962 | 0.125293 | -0.51653 | 0.154521 |
| Personal-Social DQ | -0.71834 | 0.029259 | -0.72248 | 0.027904 | -0.48395 | 0.186828 |
| Average DQ | -0.18755 | 0.628952 | -0.05192 | 0.89446 | -0.23904 | 0.535624 |
| Verbal IQ | -0.13187 | 0.487288 | -0.13132 | 0.489122 | -0.11817 | 0.533976 |
| Performance IQ | -0.22128 | 0.239948 | -0.23892 | 0.203551 | -0.29613 | 0.112077 |
| Full IQ | -0.34946 | 0.05837 | -0.3663 | 0.046494 | -0.43105 | 0.017402 |
| SRS Total | -0.0208 | 0.926805 | -0.09956 | 0.659356 | -0.07086 | 0.754014 |
| SRS Social Awareness | 0.102327 | 0.650453 | -0.01457 | 0.948698 | -0.02888 | 0.898476 |
| SRS Social Cognition | 0.169786 | 0.450013 | 0.051292 | 0.820668 | 0.013921 | 0.950973 |
| SRS Social Communication | -0.13475 | 0.549928 | -0.18957 | 0.398139 | -0.16118 | 0.473626 |
| SRS Social Motivation | 0.030698 | 0.892128 | -0.0311 | 0.89073 | 0.104288 | 0.64418 |
| SRS Autistic Mannerisms | -0.1771 | 0.430443 | -0.20464 | 0.360953 | -0.18203 | 0.417511 |

The moderate and slow developmental stages were combined as the moderate-slow stage. [a] Fasle discovery rate (FDR) correction with Benjamini-Hochberg procedure for multiple comparisons, $p$>0.05.

### Supplementary Table 10. Participants for creating group templates.

| | ASD | | | TD | | |
|---|---|---|---|---|---|---|
| | Number | Age (years, mean ± std) | Sex (% male) | Number | Age (years, mean ± std) | Sex (% male) |
| **Age 1** | 6 | 1.88±0.09 | 100 | 6 | 1.59±0.21 | 50 |
| **Age 2** | 6 | 2.61±0.32 | 50 | 6 | 2.47±0.28 | 33 |
| **Age 3** | 6 | 3.53±0.22 | 67 | 6 | 3.52±0.28 | 33 |
| **Age 4** | 6 | 4.57±0.30 | 50 | 6 | 4.65±0.29 | 50 |
| **Age 5** | 5 | 5.21±0.18 | 100 | 5 | 5.49±0.25 | 60 |
| **Age 6** | 5 | 6.69±0.35 | 100 | 5 | 6.51±0.32 | 20 |
| **Age 7** | 5 | 7.53±0.37 | 80 | 5 | 7.47±0.43 | 20 |

ASD, autism spectrum disorder. DD/ID, TD, typical developing.

**Supplementary Table 11. Smoothing parameters, EDF, $R^2$ and AIC of GAM.**

| | | TD | ASD without DD/ID | ASD with DD/ID | DD/ID |
|---|---|---|---|---|---|
| **CC Cluster** | $R^2$ (adjusted) | 0.611 | 0.567 | 0.29 | 0.204 |
| | AIC | -608.3954 | -209.1697 | -374.6034 | -168.1454 |
| | k | 6 | 4 | 4 | 3 |
| | EDF | 3.34 | 2.84 | 2.04 | 1 |
| **A-S Cluster** | $R^2$ (adjusted) | 0.741 | 0.672 | 0.489 | 0.291 |
| | AIC | -653.1693 | -207.2286 | -402.3139 | -160.5951 |
| | k | 6 | 5 | 5 | 3 |
| | EDF | 4.43 | 2.72 | 2.05 | 1 |
| **P-I Cluster** | $R^2$ (adjusted) | 0.661 | 0.613 | 0.388 | 0.206 |
| | AIC | -704.248 | -237.704 | -451.0292 | -185.0031 |
| | k | 6 | 5 | 4 | 3 |
| | EDF | 4.14 | 2.21 | 1.75 | 1 |

smoothing parameter, EDF, effective degrees of freedom. The value of EDF formed by the GAM model shows the degree of curvature of the relationship. AIC, Akaike information criterion. k, smoothing parameter, and sets the upper limit on the degrees of freedom.

### Supplementary Methods

#### Participants

All participants had no contraindications for MRI, no suspected vision or hearing problems or known genetic disorders and no neurological conditions. Children were enrolled in the ASD group if they met the DSM-5 criteria and had a diagnosis confirmed by the Autism Diagnostic Observation Schedule (ADOS). ADOS calibrated severity scores (CSSs) were calculated to allow comparisons of autism severity across participants tested by different ADOS modules. Typical development (TD) children were clinically evaluated to ensure that they had no psychiatric, neurological, or developmental disorders. TD children had an average DQ above 75 (<4 years old) or a full IQ above 70 (>=4 years old). Children with DD/ID were diagnosed by trained clinicians using the DSM-5 and had an average DQ below 75 (<4 years old) or a full IQ below 70 (>=4 years old). Children who were diagnosed before 30 months of age did not have any changes in their diagnosis in follow-up examinations.

#### Diffusion MR image quality control and exclusion criteria

The diffusion-weighted MR images of all participants were visually checked to ensure image quality in this study. We assessed the diffusion-weighted MR images carefully from the following aspects: 1) the coverage of cerebral regions (occipital/parietal/temporal/frontal lobes) was complete; 2) signal dropout due to head motion or zig-zag patterns had occurred; and 3) zipper artefacts[1], the most common equipment artefacts, were caused by radio frequency (RF) break-through (see Supplementary Fig. 1). Images were excluded if they met the following exclusion criteria: 1) more than 3 volumes (more than 5% of all volumes) that exhibited signal dropout, zig-zag patterns and/or zipper artefacts; and 2) incomplete coverage of the 36 selected core white matter ROIs. Based on these criteria, a total of 40 participants were excluded, including 16 ASD with DD/ID, 5 ASD without DD/ID, 17 TD and 2 DD/ID children.

#### Participants for creating group templates

Seven study-specific and age-specific tensor templates were created from tensor maps of a selective subset of participants within the same year. We selected 5 to 6 children in the ASD subgroups and TD group per year with excellent image quality (see Supplementary Table 10).

#### Cubic splines

The estimated nonparametric functions *s(age)* were obtained using cubic splines in R version 4.2.1. In the cubic spline function, the entire age range was divided into 100 intervals of equal size, and each interval was represented by a cubic equation. *s(age)* can be represented using the following formula:

$$s(age)=a*(x_i - x_0)^3+b*(x_i - x_0)^2+c*(x_i - x_0)+d,$$

where $x_i$ represents the age of the $i^{th}$ subject, and $x_0$ represents the starting age of the interval to which the $i^{th}$ subject belongs.

#### Group comparisons and stage comparisons using general linear model

For each white matter cluster, we used a general linear model (GLM) to evaluate group differences in both the NDI and its growth rates in various stages of early childhood. Notably, the moderate and slow developmental stages were combined into a moderate-slow stage due to their short durations. Age, sex, and head motion during the MRI scan were included as covariates to account for potential confounding effects. Head motion was quantified by the AMzscore and RMzscore, referring to the z score of the root-mean-square displacement of the mean absolute intervolume displacement and mean relative intervolume displacement, respectively.

To compare the group differences in the NDI within each white matter cluster, we constructed the GLM using the following formula:
1) NDI ~ age + sex + AMzscore + RMzscore + groups.
To compare the group differences in the growth rate within each white matter cluster, we constructed the GLM with the following formula:
2) Growth rate~ age + sex + AMzscore + RMzscore + groups.
To compare the inter-stage differences in the NDI within each white matter cluster and each group, we constructed the GLM with the following formula:
3) NDI ~ age + sex + AMzscore + RMzscore + stages.
False discovery rate (FDR) correction using the Benjamini–Hochberg procedure was conducted for multiple comparisons.
